# The blood-brain barrier regulates brain tumour growth specifically via the SLC36 amino acid transporter Pathetic in *Drosophila*

**DOI:** 10.1101/2025.05.15.654179

**Authors:** Qian Dong, Edel Alvarez-Ochoa, Hina Kosakamoto, Fumiaki Obata, Cyrille Alexandre, Louise Cheng

## Abstract

Tumours adapt their metabolism to sustain increased proliferation, rendering them particularly vulnerable to fluctuations in nutrient availability. However, the role of the tumour microenvironment in modulating sensitivity to nutrient restriction (NR) remains poorly understood. Using a *Drosophila* brain dedifferentiation neural stem cell (NSC) tumour model induced by Prospero (Pros) inhibition, we show that tumour sensitivity to NR is governed by the perineural glial (PG) cells of the blood-brain barrier (BBB), a major component of the glial niche surrounding the tumour. We identify the SLC36 amino acid transporter Pathetic (Path) as a crucial regulator of nutrient sensitivity. Under NR, while wildtype buffers against low nutrient levels by upregulating Path, tumour glia downregulate Path. Furthermore, Path is specifically required by the tumour (but not wildtype) PG; its downregulation causes reduced proliferation of PG cells and, in turn, restricts NSC tumour growth. Path influences PG proliferation via the mTor-S6K pathway, and its expression is controlled by Ilp6 levels and the Insulin/PI3K pathway. Overexpression of Path is sufficient to counteract the inhibitory effects of NR on tumour growth. These findings suggest that Path levels at the BBB play a key role in determining tumour sensitivity to NR.

## Introduction

Altered metabolism is a hallmark of cancer [1]. In particular, disruptions in amino acid metabolism have been shown to selectively inhibit tumour growth based on their genetic profile and tissue of origin [2, 3]. Identifying the metabolic vulnerabilities linked to these cancer-specific characteristics offers opportunities for novel dietary and therapeutic interventions targeting cancer growth. In brain tumours, moderate dietary restriction has been shown to significantly reduce astrocytoma growth [4], while cysteine and methionine withdrawal sensitise gliomas to ferroptosis [5]. Additionally, elevated glucose levels are associated with increased proliferation and poor drug response in glioblastoma cell lines [6, 7]. Despite the therapeutic potential of dietary interventions in brain cancer, the mechanisms linking diet to brain tumour growth remain unclear, highlighting the need for further investigation into the relationship between metabolism and tumour progression.

In this study, we utilise a *Drosophila* brain tumour model induced via dedifferentiation, a central mechanism underlying tumour formation, to study the effect of nutrient restriction (NR) on tumorigenesis. *Drosophila* has a simple central nervous system (CNS) where neural stem cells (NSCs), called neuroblasts (NBs), undergo asymmetric divisions to produce diverse neuronal and glial populations that form adult fly brains (Fig 1A). During brain development, NBs reside in a specialised microenvironment, or niche, which provides essential signals and trophic support to regulate NB behaviour [8, 9]. This niche consists of cortex glial (CG) cells, which form individual chambers to house each NB and its progeny [10–14]. Additionally, the surface glial cells, including the perineural glia (PG) and sub-perineural glia (SPG), collectively form the blood-brain barrier (BBB) at the brain surface, which is covered by a sheath of extracellular matrix known as the neural lamella (Fig 1C) [15–17]. This barrier prevents the passage of drugs or toxins while facilitating the uptake of nutrients via amino acid, glucose, and lipid transporters [16, 18–21].

**Fig 1.**
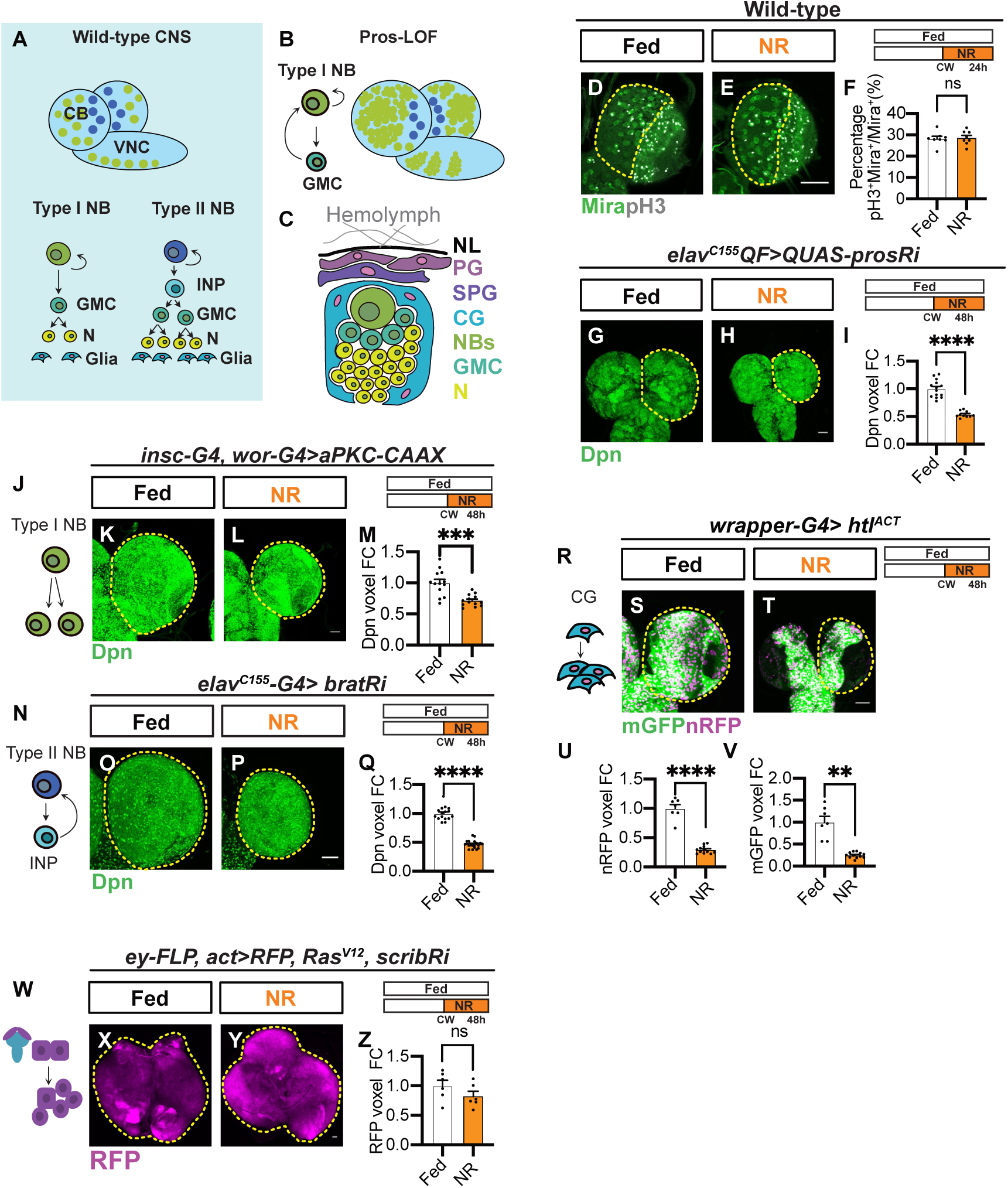
The growth of brain tumours is sensitive to post-critical weight (CW) nutrient restriction (NR). (A) Schematic representation of the larval central nervous system (CNS). The CNS consists of two brain lobes and one ventral nerve cord (VNC). Type I neuroblasts (NBs, green), in the central brain (CB) region of brain lobes and VNC, undergo asymmetric divisions to produce a self-renewing NB and a ganglion mother cell (GMC), which further divides once to make two differentiated neurons or glia (yellow). Type II NBs (blue) are limited to a specific region on the dorsal side of the CB. They divide asymmetrically like Type I but generate an intermediate neural progenitor (INP), which undergoes several rounds of division, producing multiple GMCs and, thus, more neurons or glia than type I NBs. (B) Schematic depicting the formation of Prospero (Pros) Loss-of-function (LOF) NB tumours. Loss of the transcription factor Pros reverses GMCs to type I NBs (green), which go through unlimited divisions to form NB tumours. (C) Schematic representation of the NB niche, which is composed of perineural glia (PG) and sub-perineural glia (SPG), forming the blood-brain barrier (BBB) at the brain surface under neural lamina (NL); and cortex glia (CG), which directly encase NBs, GMCs and neurons (Ns). (D-E) Single-section images of wild-type brain lobes (ventral side) stained with the NB marker Miranda (Mira) and the mitotic marker pH3 under Fed and NR conditions. CBs are circled by yellow dashed lines. NR: 63.5-96hALH; Dissection: 96hALH. Genotype: *repo-G4>UAS-RFP*. (F) Quantification of the percentage of type I NBs undergoing mitosis (pH3^+^Mira^+^) in the CB (D-E) (n = 8, 8). (G-H) Maximum projection images of *elav^C155^QF>QUAS-prosRi* tumour brains stained with the NB marker Deadpan (Dpn) under Fed and NR conditions. (I) Quantification of normalised (to Fed) Dpn voxels in (G-H) (n = 14, 10). (J-L) Overexpression of *aPKC-CAAX* with the NB driver *insc-G4, wor- G4* induces symmetric division of type I NB (J) and formation of NB tumours. (K-L) are maximum projection images of *insc-G4, wor-G4> aPKC-CAAX* tumour brain lobes marked by Dpn under Fed and NR conditions. (M) Quantification of normalised (to Fed) Dpn voxels of circled brain lobes in (K-L) (n = 13, 13). (N-P) Overexpression of *bratRNAi* with the NB driver *elav^C155^-G4* reverts INPs to type II NBs (N), forming NB tumours in the dorsal side of the CB. (O-P) are maximum projection images of *elav^C155^-G4*> *bratRNAi* tumour brain lobes marked by Dpn under Fed and NR conditions. (Q) Quantification of normalised (to Fed) Dpn voxels of circled brain lobes in (O-P) (n = 16, 20). (R-T) Overexpression of *htl^ACT^* with the CG driver *wrapper-G4* causes over-proliferation of CG (R). (S-T) are maximum projection images of *wrapper-G4> htl^ACT^* brain lobes marked with mGFP and nRFP under Fed and NR conditions. (U-V) Quantification of normalised (to Fed) nRFP voxels and mGFP voxels of circled brain lobes in (S-T), respectively (n = 7, 12). (W-Y) Co-expression of *Ras^V12^*and *scribRNAi* in eye discs (*ey-FLP, act-G4*) leads to the formation of neoplastic tumours (marked by *UAS-RFP* and circled in X-Y). (X-Y) are maximum projection images of eye disc tumours under Fed and NR conditions. (Z) Quantification of normalised (to Fed) RFP voxels in (X-Y) (n = 6, 6). Data information: ALH = after larvae hatching. NR: 72-120hALH; Dissection: 120hALH unless otherwise stated. Scale bar = 50μm. Error bar represents SEM. In (F): unpaired t-test, (ns) P = 0.8060. In (I): Welch’s t-test, (****) P < 0.0001. In (M): Welch’s t-test, (***) P = 0.0003. In (Q): Welch’s t-test, (****) P < 0.0001. In (U): Welch’s t-test, (****) P < 0.0001. In (V): Welch’s t-test, (**) P = 0.0013. In (Z): unpaired t-test, (ns) P = 0.2013.

Type I NBs are the most common NB type found throughout the CNS (Fig 1A). They produce neurons by first generating an intermediate progenitor cell called ganglion mother cell (GMC), where the homeodomain transcription factor Prospero (Pros) promotes its differentiation into two postmitotic neurons [22]. We previously showed that NB lineages can buffer against nutrient shortage, and the ability to do so is determined by the activation of the PI3K signalling pathway, via the glial niche secreted ligand Jelly Belly (Jeb) and NB-specific Anaplastic Lymphoma Kinase (ALK) [24–26]. Here, we found that the brain tumours caused by the loss of Pros are in fact sensitive to nutrient restriction (NR) (Fig 1B) [23]. This sensitivity is attributed to the downregulation of the SLC36A4 amino acid transporter, Pathetic (Path) under NR, which leads to reduced PG proliferation and, in turn, tumour growth. In contrast, Path is upregulated in control brains upon NR and is not required under fed conditions. Path influences PG proliferation via the mTor-S6K pathway, and its expression is controlled by Ilp6 levels and the insulin/PI3K pathway. Notably, overexpressing Path is sufficient to counteract the inhibitory effects of NR on tumour growth. Surprisingly, Path does not facilitate the transport of its known substrates – proline, alanine, or tryptophan – into the brain. Instead, it regulates the availability of amino acids, including branched-chain amino acids (BCAAs), which are critical for PG proliferation and tumour growth. These findings suggest that Path levels at the BBB play a key role in determining the response of brain tumours to NR.

## Results

### The growth of dedifferentiation-induced NB tumours is sensitive to NR

We previously demonstrated that neurogenesis becomes insensitive to dietary restriction after critical weight (CW, around 60 hours after larval hatching, ALH), a developmental checkpoint beyond which starvation no longer impacts the initiation of metamorphosis or survival [24–26]. Consistent with this, in wild-type brains, we found that both NB number and cell cycle progression remained unaffected, following 24 hours of nutrient restriction (NR) post-CW (NR: complete starvation on 0.4-1% agar, S1A and I Fig, Fig 1D-F, pH3 index). Next, we assessed whether, under stressful conditions, such as in the case of brain tumours, the brain would respond differently. We induced dedifferentiation-derived NB tumours by expressing *QUAS-prosRNAi* using the pan-NB lineage driver, *elav^C155^-QF2*, and subjected the animals to NR starting from CW. Brain tumours in *Drosophila* induced by *pros* knockdown cause developmental delays due to the disruption of ecdysone signalling (as shown by [27]). To investigate the relationship between tumour growth and NR, we first identified when tumour-bearing larvae reached CW. By subjecting these larvae to NR at various developmental stages, we assessed the temporal sensitivity of tumours to nutrient availability (S1A Fig). We determined that tumour-bearing *Drosophila* larvae reached CW at approximately 68 hours after larval hatching (ALH). Larvae subjected to NR before this time point exhibited severe developmental consequences: those starved from 48 hours ALH died and those starved from 63.5 hours ALH failed to pupariate on time (S1A-B Fig). In comparison, subjecting the animals to NR from 68 hours ALH and onwards caused them to pupariate earlier (S1A-B Fig) [28]. NR effectively suppressed the growth of most polyploid tissues in tumor-bearing animals, including the fat body and salivary glands (S1C-D and F-G Fig), similar to its effects in wild-type animals [24]. Unlike wild-type brains, brain tumours exhibited significant growth reduction following 24 or 48 hours of NR post-CW compared to fed conditions (S1E and H Fig, Fig 1G-I, S2A-C Fig). From this point onward, tumour-bearing animals were subjected to NR starting at 72 hours ALH, either for 24 hours when compared with wildtype controls (which reach the wandering stage 24 hours after CW), or for 48 hours unless otherwise specified. These findings highlight the unique sensitivity of brain tumours to NR, even when normal brain proliferation is unaffected.

To elucidate the mechanisms accounting for the reduced tumour growth, we examined tumour proliferation, differentiation and cell death. The reduction in brain tumour size following NR was found to be due to a slowdown in cell cycle progression. This was evidenced by a decrease in the incorporation of the thymidine analogue EdU (5-ethynyl-2′- deoxyuridine) into the S phase of the cell cycle during a 15-minute labelling window (S2D-F Fig). As the tumour arises from dedifferentiation, we next assessed whether the tumour size reduction upon NR is due to increased differentiation from NBs into neurons. We observed a slight decrease in the percentage of neurons (Elav/DAPI ratio, S2G-I Fig), indicating that NR reduces tumour growth not by inducing more conversion of NBs to neurons. The reduction in brain tumour size following NR was also not accounted for by increased cell death, as indicated by the unaltered percentage of NBs marked by the apoptotic marker Dcp-1 (S2J-L Fig). This suggests that NR impedes tumour growth primarily by slowing down cell cycle progression, rather than by promoting differentiation or inducing apoptosis.

We next assessed whether NR affects the growth of different tumour types, including other brain and epithelial tumours. We found that NR significantly reduced the size of brain tumours induced in (1) type I NBs by the constitutive activation of aPKC (Fig 1J) [29]; (2) type II NBs by the knockdown of the NHL family protein Brat (Fig 1N) [30]; and (3) cortex glial (CG) cells, by the constitutive activation of the FGF receptor Heartless (Fig 1R) [31]. In all cases, tumour size was significantly reduced after 48 hours of NR (Fig 1K-M, O-Q, and S-V). These findings demonstrate that NR broadly impacts tumour growth across various brain tumour models. In contrast, the growth of the eye-antennal disc tumours, induced via the activation of Ras (*Ras^V12^*) and the knockdown of Scribble (*scrib RNAi*) (Fig 1W) [32], was not significantly affected by NR (Fig 1X-Z). Therefore, NR sensitivity could represent a distinctive characteristic of brain tumours, distinguishing them from wild-type tissue and epithelial tumours.

### The BBB PG glial expansion is significantly reduced under NR in tumour brains

The brain tumour lineages are surrounded by a glial niche made up of CG, PG and SPG (Fig 1B-C) [8]. This glial niche has been shown to provide the trophic support necessary for wild-type neural proliferation [8]. These glial cells likely play a key role in modulating the tumour microenvironment and supporting tumour growth, which may contribute to the observed sensitivity of brain tumours to NR. To investigate whether the glial niche regulates the response of NB tumours to NR, we first examined how the number of glial cells, marked by the pan-glial nuclear marker Reversed Polarity (Repo), is altered under NR conditions. We found that the overall number of glial cells was reduced by 35.25% after 24 hours of NR and 47.78% after 48 hours of NR compared to Fed (Fig 2A-F). Similarly, we found that *bratRNAi* and *aPKC-CAAX* brain tumours also exhibited significant reductions in glial cell number upon 48 hours of NR compared to Fed (S3A-F Fig). In contrast, wild-type brains exhibited a glial number reduction by only 15.65% after 24 hours of NR. (Fig 2G-I) [33], suggesting that NR has a greater effect on glial numbers in tumour brains than in wild-type brains.

**Fig 2.**
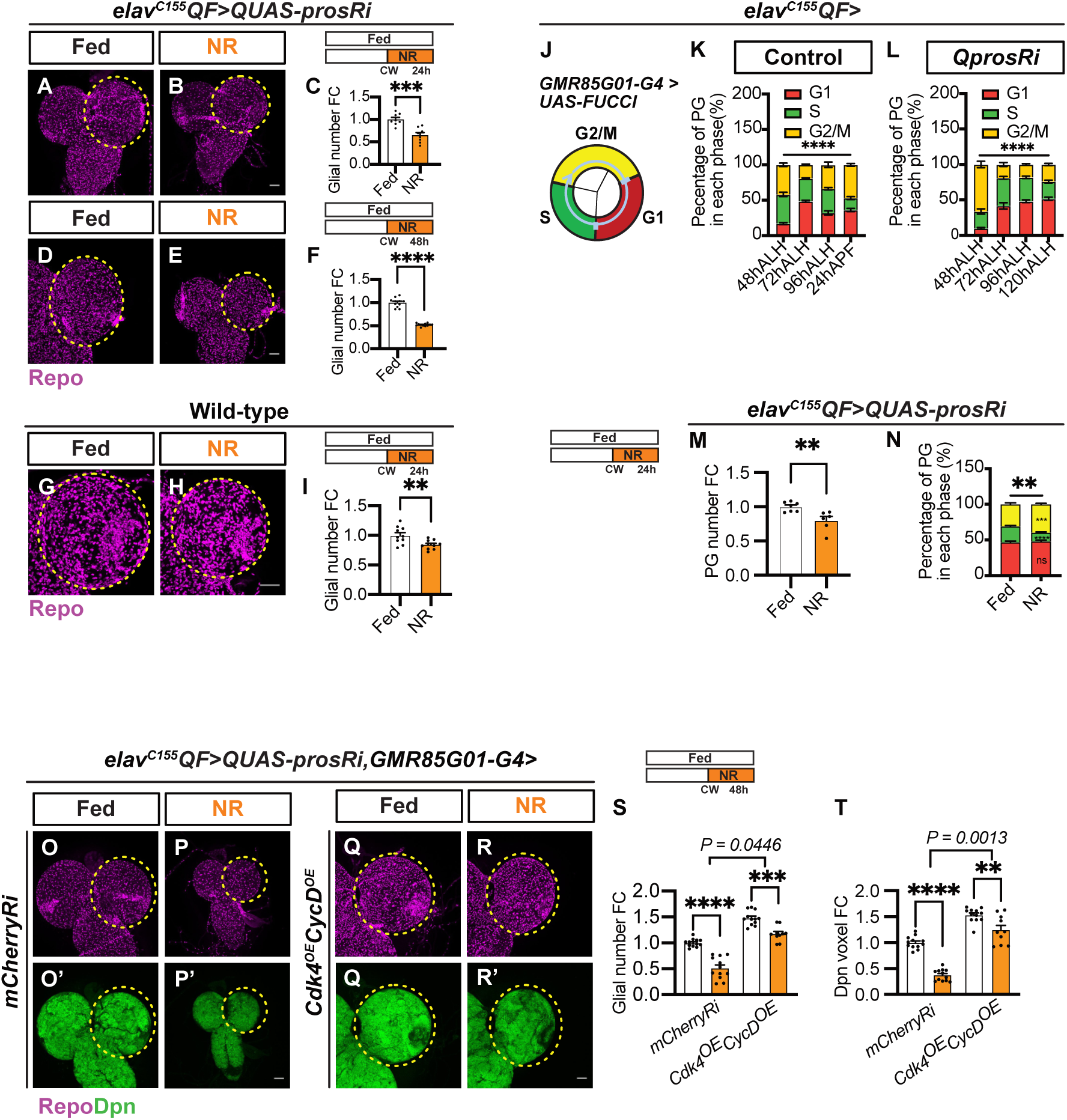
Blood-brain barrier (BBB) perineural glia (PG) proliferation is compromised under nutrient restriction (NR), and this underlies the high susceptibility of brain tumours to NR. (A-B) Maximum projection images of *elav^C155^QF>QUAS-prosRi* tumour brains stained with the pan-glial marker Repo under Fed vs NR. NR: 72-96hALH; Dissection: 96hALH. (C) Quantification of normalised (to Fed) glial number in the brain lobe in (A-B) (n = 8, 8). (D-E) Maximum projection images of *elav^C155^QF>QUAS-prosRi* tumour brains stained with the pan-glial marker Repo under Fed vs NR. NR: 72-120hALH; Dissection: 120hALH. (F) Quantification of normalised (to Fed) glial number in the brain lobe in (D-E) (n = 11, 12). (G- H) Maximum projection images of wild-type brain lobes stained with Repo under Fed vs NR. NR: 63.5-96hALH; Dissection: 96hALH. (I) Quantification of normalised (to Fed) glial number in the brain lobe in (G-H) (n = 10, 10). (J) Schematic depicting Fly-FUCCI, a tool that enables the visualisation of cell cycle transitions by tagging E2F1 and Cyclin B proteins with ECFP (red) and Venus (green). As the cell cycle progresses, these proteins are targeted for degradation by the S phase-dependent ubiquitin ligase CRL4Cdt2 and the ubiquitin E3- ligases APC/C during mitosis, respectively. When FUCCI is overexpressed using the *GMR85G01-G4*, it allows the assessment of PG cell cycle progression in (K-L and N) and total number in (M). (K-L) Quantification of the percentage of PG cells in each cell cycle phase within the circled brain lobes in (Fig. S3G-H’’’) (n = 10, 7, 7, 4, 8, 5, 6, 4). (M-N) Quantification of the total number of PGs (M) and the percentage of PG cells in each cell cycle phase (N), labelled by FUCCI using *GMR85G01-G4* in each *elav^C155^QF>QUAS-prosRi* tumour brain lobe under Fed vs NR (n= 7, 6). NR: 72-96hALH; Dissection: 96hALH. (O-R’) Maximum projection images of *elav^C155^QF>QUAS-prosRi* tumour brains where *Cdk4* and *CycD* were co-expressed in PG using *GMR85G01-G4* in (Q-R’), compared with *mCherryRi* (O-P’) under Fed vs NR. Glial cells are marked with Repo, and NB tumours are labelled with Dpn. NR: 72-120hALH; Dissection: 120hALH. (S-T) Quantifications of the normalised (to *mCherryRi* fed condition) glial number (S) and Dpn voxels (T) of each circled brain lobe in (S-T’) (n = 12, 12, 12, 10). Data information: ALH = after larvae hatching. APF = after pupa formation. Brain lobes are circled with yellow dashed lines. Scale bar = 50μm. Error bar represents SEM. In (C): unpaired t-test, (***) P = 0.0003. In (F): Welch’s t-test, (****) P < 0.0001. In (I): unpaired t- test, (**) P = 0.0072. In (K and L): Two-way ANOVA were used to analyse the distribution of PG cells in each cell cycle phase over time in both wild-type brains and tumour-bearing brains. In (K): interaction P < 0.0001. In (L): interaction P < 0.0001. Mean± SEM, and statistical results including multiple comparisons are displayed in S1A-B Table. In (M): unpaired t-test, (**) P = 0.0073. In (N): Chi-square test, (**) P = 0.0014. Mean± SEM, and statistical results, including two-way ANOVA multiple comparisons, are displayed in S1C Table. In (S-T): Two-way ANOVA were used to analyse whether the effect of NR on glial number and tumour size is affected by *CdK4* and *CycD* overexpression (significance indicated by interaction P value). In (S): interaction P = 0.0446; In (T): interaction P = 0.0013. Mean± SEM, and statistical results including multiple comparisons are displayed in S1D-E Table.

The BBB, composed of PG and SPG, regulates nutrient transport into the brain. Next, we investigated how the BBB glia were affected by NR. During development, SPG accounts for a minority of all glial cells [15–18]. Therefore, the observed change in glial cell number under NR is unlikely to be due to alterations in SPG cell numbers. Instead, PG cells proliferate through mitosis, rather than undergoing endoreplication, during neurogenesis. [15, 31]. Using the Fly-FUCCI tool [34] driven by the PG-specific *GMR85G01-GAL4* driver [35] (Fig 2J), we observed that PG cells actively progressed through the cell cycle during early larval stages and continued to proliferate post-CW in wild-type and tumour brains (S3G-H’’’ Fig and Fig 2K-L). In tumour-bearing brains, however, PG cell cycle slowed progressively over time, as indicated by an increase in the percentage of G1-phase cells (S3H-H’’’ Fig and Fig 2L).

Upon subjecting tumour-bearing larvae to 24 hours of NR, we found that the total number of PG cells in the BBB was significantly reduced compared to fed controls (Fig 2M), suggesting that NR impacts PG proliferation and may contribute to tumour growth inhibition. We examined FUCCI to assess if this reduction in PG number is accounted for by their slower proliferation. We showed that the number of PG cells in the S phase was significantly decreased, and those in the G2/M phase were significantly increased after 24 hours of NR (Fig 2N), indicating that the cell cycle progression of PG cells is further reduced. Increased G2/M cell cycle arrest is typically associated with increased cell death [36], but we did not observe a significant change in glial cell death following NR (S3I-K Fig). These findings suggest that NR impacts PG cell cycle progression, leading to a decrease in PG numbers and possibly tumour growth inhibition as a consequence.

To evaluate whether enhancing PG cell proliferation could promote tumour growth under NR, we overexpressed the cell cycle genes *Cdk4* and *CycD,* specifically in PG cells, using the *GMR85G01-GAL4* driver. Compared to the control (*mCherryRNAi*), the overexpression of *Cdk4* and *CycD* increased overall glial number under fed conditions (Fig 2Q, O and S). Furthermore, this manipulation partially rescued the decrease in glial numbers caused by NR (Fig 2O-R and S, the number of glia is reduced 48% upon NR in *mCherryRi*, and 20% in *Cdk4^OE^, CycD^OE^*, statistical assessment described in Methods). As a result, the decrease in tumour size upon NR was also partially rescued in the *Cdk4/CycD*- overexpressing condition compared to *mCherryRNAi* control (Fig 2O’-R’ and T, the size of tumour is reduced 63% upon NR in *mCherryRi*, and 18% in *Cdk4^OE^, CycD^OE^*, statistical assessment described in Methods). These findings suggest that manipulating PG cell numbers can influence the sensitivity of brain tumour growth to NR, highlighting the role of PG proliferation in tumour response to nutrient availability.

### Branched-chain amino acids (BCAAs) are required for glial and tumour expansion

During development, growth is fuelled by nutrients derived from the diet, including carbohydrates and proteins. To identify the essential nutrient components for glial and tumour expansion, we selectively removed yeast (protein source) or glucose and polenta (sugar source) from the standard diet starting at 48 hours ALH. We then assessed the impact of these nutrient restrictions on glial and tumour growth, aiming to determine which nutrients are critical for supporting the proliferation of glial cells and tumour progression. We found that the withdrawal of yeast as well as glucose and polenta, reduced glial number and tumour growth, phenocopying the effects of NR (Fig 3A-C and F-G). Notably, the inhibitory effect was more pronounced when tumour-bearing larvae were placed on a yeast-free diet (Fig 3A- C and F-G). To test the sufficiency of yeast in promoting tumour growth, we supplemented the diet with 2x yeast from 48 hours ALH. We found that increasing yeast content modestly increased both glial cell numbers and tumour size (Fig 3D-E, and F-G).

**Fig 3.**
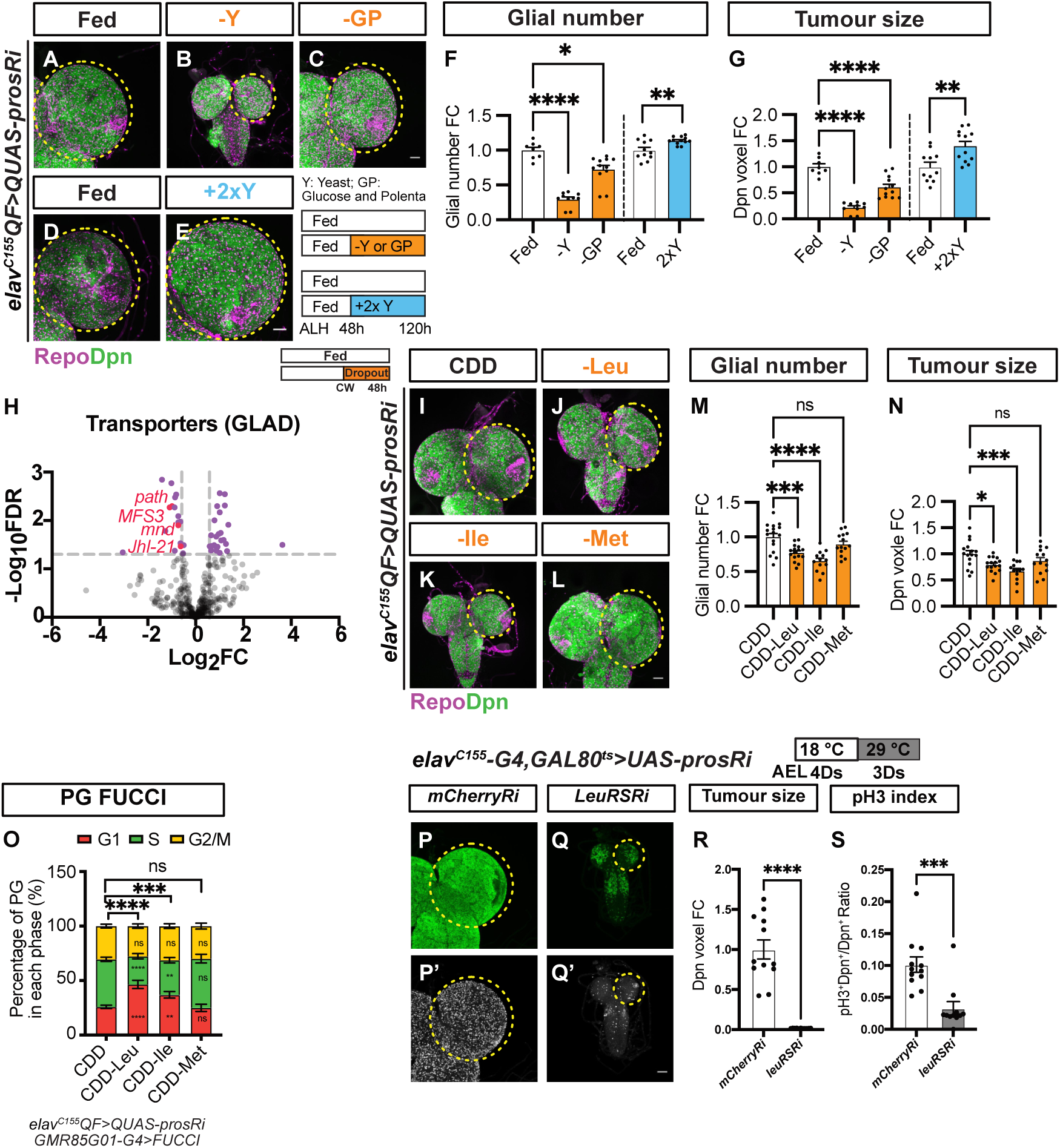
Branched-chain amino acids (BCAAs) are necessary for glial and neuroblast (NB) tumour expansion. (A-E) Maximum projection images of *elav^C155^QF>QUAS-prosRi* tumour brains stained with the glial marker Repo and NB marker Dpn, under fed (A), dropout of Yeast (B) or glucose and polenta (C) from 48-120hALH; and fed (D) vs 2xYeast supplementation (E) from 48-120hALH. (F-G) Quantifications of the normalised (to Fed) glial number (R) and Dpn voxels (S) of each circled brain lobe in (A-E) (n = 8, 10, 12, 11, 12). (H) Volcano Plot depicting differential gene expression in the *elav^C155^QF>QUAS-prosRi* tumour brains under NR compared to Fed. Genes that are significantly altered (False discovery rate FDR <0.05; FC > 1.5) are marked with purple dots. NR: 72-120hALH; Dissection: 120hALH. (I-L) Maximum projection images of *elav^C155^QF>QUAS-prosRi* tumour brains stained with the glial marker Repo and NB marker Dpn. The animals were moved from standard food to CDD (I), CDD- Leu (J), CDD-Ile (K) and CDD-Met (L) at 72hALH and dissected at 120hALH. CDD: chemically defined diet. (M-N) Quantifications of the normalised (to Fed) glial number (M) and Dpn voxels (N) of each circled brain lobe in (I-L) (n = 16, 16, 13, 14). (O) Quantifications of PG cell cycle distribution in *elav^C155^QF>QUAS-prosRi* tumour brain lobes, where FUCCI was overexpressed in PG using *GMR85G01-G4*, under fed (CDD), dropout of Leu, Ile or Met (n = 10, 5, 8, 6). (P-Q) Maximum projection images of *elav^C155^- G4, GAL80^ts^>UAS-prosRi* tumour brains with *mCherryRi* or *LeuRSRi* overexpression. Larvae were placed at 18°C for 4 days before being moved to 29°C for transgene activation (3 days). (P and Q): Dpn; (P’ and Q’): phosphorylated histone H3 (pH3). (R-S) Quantifications of the normalised (to *mCherryRi*) Dpn voxels (R) and pH3 index (S) of each circled brain lobe in (P-Q) (n = 12,11) Data information: ALH = after larvae hatching. Brain lobes are circled with yellow dashed lines. Scale bar = 50μm. Error bar represents SEM. In (F): Kruskal-Wallis test, (****) P < 0.0001; (*) P = 0.0463; Welch’s t-test, (**) P = 0.0089. In (G): One-way ANOVA, (****) P < 0.0001; (****) P < 0.0001; unpaired t-test, (**) P = 0.0040. In (M): One-way ANOVA, (***) P = 0.0006; (****) P < 0.0001; (ns) P = 0.2437. In (N): One-way ANOVA, (*) P = 0.0189; (***) P = 0.0002; (ns) P = 0.2308. In (O), the Chi-square test was used to compare the difference in cell cycle phase distribution between nutritional conditions. (****) P < 0.0001; (***) P = 0.0001; (ns) P = 0.6442. Two-way ANOVA was used to compare the differences in each cell cycle phase between nutritional conditions. Mean± SEM, and statistical results, including multiple comparisons, are displayed in S1F Table. In (R): Welch’s t-test, (****) P < 0.0001. In (S): Mann Whitney test, (***) P = 0.0004.

Amino acids constitute a significant component of yeast. Using Liquid Chromatography-Tandem Mass Spectrometry (LC-MS/MS), we investigated the impact of NR on amino acid (AA) levels. Our analysis revealed a substantial reduction in AA levels in both the haemolymph and the tumour brains (S4A-B Fig). AAs are imported into the brain through transporters located on its surface. To further understand how NR affects the brain’s nutrient uptake, we examined the expression of nutrient transporters in tumour brains under NR conditions. We compared the expression of annotated transporters (using Gene List Annotation for *Drosophila* (GLAD)) in tumour brains under fed and NR conditions via bulk RNA-sequencing. We showed that the gene expression profile of tumour brains under NR is separated from that of control-fed tumour brains (S5A-B Fig), with 225 genes significantly upregulated and 301 significantly downregulated (False Discovery Rate (FDR) ≤0.05, Fold Change (FC) ≥ 1.5, S1 Data). We found that AA transporters, including Minidiscs (MND), Juvenile hormone Inducible-21 (JhI-21) and Pathetic (Path), were among the top transcriptionally downregulated transporters in the tumour brain under NR (Fig 3H).

Furthermore, we observed a reduction in the carbohydrate transporter Major Facilitator Superfamily Transporter 3 (MFS3), the lipid transporter fatty acid binding protein (FABP) and several ATP synthetase components and subunits (S2 Data).

MND and JhI-21 are reported to be transporters for branched-chain amino acids (BCAAs), i.e. leucine (Leu), isoleucine (Ile) and valine (Val) [37–39]. Given that both *mnd* and *Jhl-21* were transcriptionally downregulated upon NR, we next tested whether BCAAs are rate-limiting for the expansion of glia and tumours, using a chemically defined diet (CDD) [40, 41]. We showed that the removal of Leu or Ile from 72 hours ALH resulted in a reduction in glial cell number and tumour size (Fig 3I-N). Brain tumours are known to be addicted to dietary methionine (Met) [42]. However, in our study, the withdrawal of Met did not significantly affect glial cell number or tumour growth (Fig 3I-N). Our data indicate that BCAAs are crucial dietary components for glial and tumour growth. To further investigate their role, we analysed the cell cycle profiles of PG cells under Leu or Ile dropout conditions. We found that the reduction in glial numbers was driven by a slowdown in PG cell cycle progression, as evidenced by a significant increase in the percentage of cells in the G1-phase and a significant reduction in the number of cells in the S-phase (Fig 3O). In comparison, these changes were not observed under Met dropout (Fig 3O). These results suggest that BCAAs play a key role in regulating PG proliferation and, consequently, tumour growth.

We next examined how deprivation of BCAAs slows down tumour growth. BCAAs particularly Leu, are known to support cell growth by promoting protein synthesis and by fueling the Tricarboxylic Acid (TCA) cycle [43]. In line with this, our RNA-seq data showed that the Aminoacyl-tRNA biosynthesis and the TCA cycle are among the top 10 downregulated pathways under NR compared to fed conditions (S5C Fig). We found that leucyl-tRNA synthetase, isoleucyl- tRNA synthetase, but not valyl- tRNA synthetase, which function to conjugate the amino acids to the corresponding tRNA for protein synthesis at the ribosome, are significantly downregulated under NR (S5D Fig). However, enzymes including branched-chain amino acid transaminase (BCAT) and branched-chain alpha-ketoacid dehydrogenase complexes (Bckdha, Bckdhb and Dbct, identified and classified in flies in [44]), which convert BCAAs to the TCA cycle, are not altered under NR (S5D Fig). These indicate that downregulation of BCAA tRNA synthetases might account for why NB tumour growth is compromised under NR. Indeed, temporal knockdown of leucyl-tRNA synthetase in the NB tumour (*elav^C155^-GAL4> prosRi*) dramatically reduced the tumour size and the percentage of tumour cells that are pH3^+^(pH3 index, Fig 3P-S), suggesting that leucyl-tRNA synthetase is necessary for NB tumour growth. Similarly, this manipulation also slows down wild-type brain growth (DAPI staining, S5E-G Fig), suggesting the role of leucyl-tRNA synthetase during brain development. However, we found that in wild-type animals, post-CW NR did not change the expression level of this enzyme (S5H Fig). This partially underlies why the wild-type brains are spared under post-CW NR. Collectively, our data suggest that dietary BCAAs are required for brain tumour growth via two mechanisms: first, they regulate the expansion of PG, which controls the amount of BCAAs that can be transported into the brain; and second, they promote protein synthesis of NB tumours to support their growth.

### The amino acid transporter Pathetic (Path) is downregulated in PG of tumour brains under NR

RNA sequencing data indicated that the expression of the SLC36 amino acid transporter Pathetic (Path) is significantly downregulated in tumour brains under NR. To further investigate this, we examined changes in Path protein expression using a Path-GFP protein trap [45]. Consistent with previous studies, we observed that Path-GFP was expressed in glial cells, marked by *repo-GAL4>mRFP* (S6A-C Fig) [25], and NB tumours marked by Mira (S6G-H Fig). We showed that its expression at the surface glia of the brain was comparable between wild-type and tumour brains (S6A-C Fig). Under NR, in wild-type brains, Path-GFP levels were elevated in the surface glia (S6D-F Fig). In contrast, Path-GFP expression in the surface glia of tumour brains was significantly reduced compared to fed conditions (Fig 4A-D). This finding suggests that NR induces differential regulation of Path expression in glial cells depending on the presence of tumours. In addition, we observed a reduction in Path-GFP expression in both CG cells (the glia that form a chamber surrounding NB tumours) and tumour NBs under NR (S6G-I Fig). Interestingly, NR-resistant *Ras^V12^scrib^RNAi^* tumour exhibited no significant change to the Path expression level in response to NR (S6J-L Fig). Collectively, these findings suggest that Path expression may correlate with tumour sensitivity to NR.

**Fig 4.**
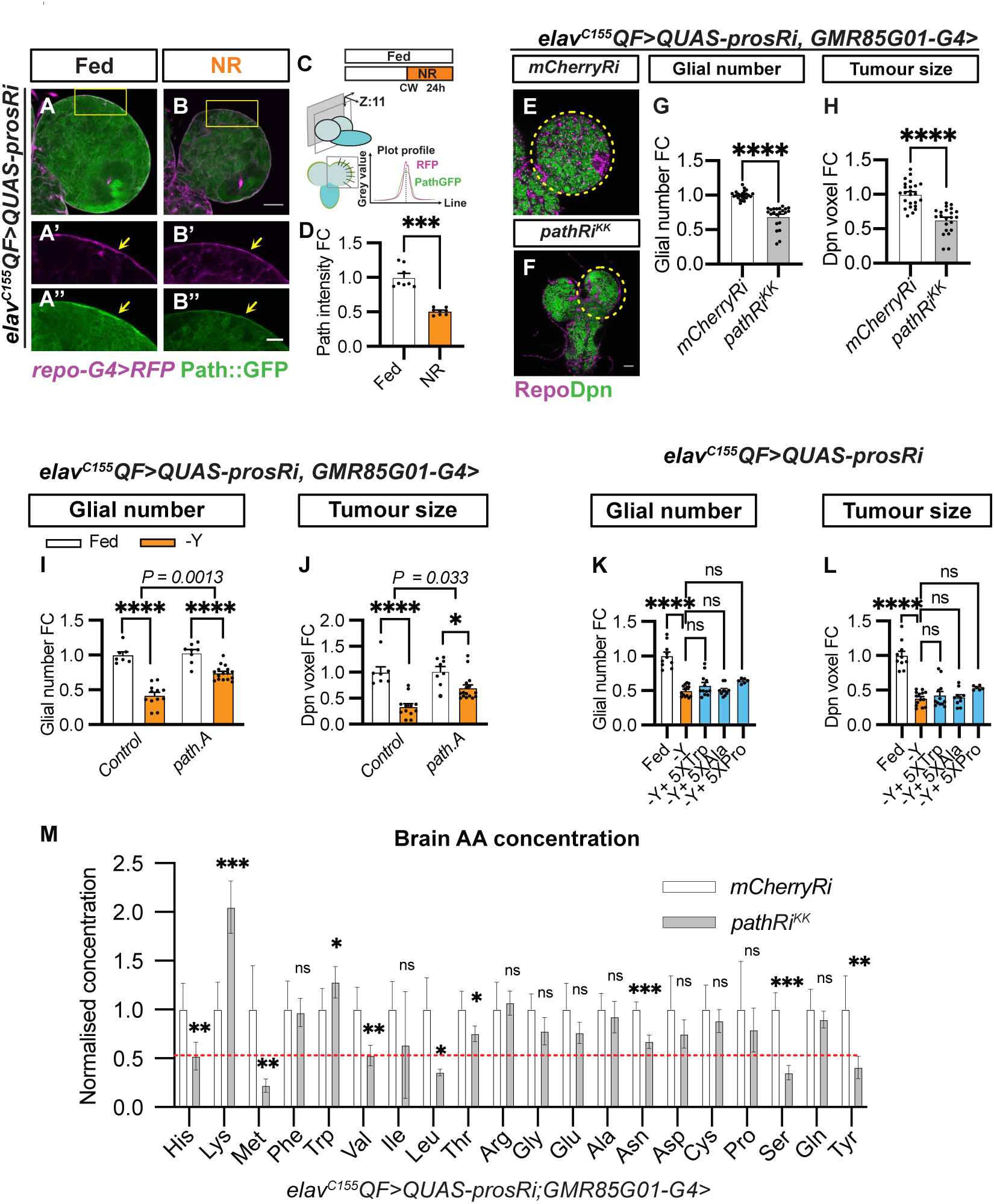
Pathetic (Path) is downregulated at the blood-brain barrier (BBB) under nutrient restriction (NR), restricting the availability of essential amino acids (EAAs) for tumour growth. (A-B) Single-section (A-B) and zoomed-in images (A’-B’’) of *elav^C155^QF>QUAS-prosRi* tumour brains where Path-GFP is expressed in the glia (marked by *repo-G4>mRFP*) at the brain surface and is downregulated under NR compared to Fed conditions. NR: 72-96hALH; Dissection: 96hALH. Scale bar = 20μm in (A’-B’’) (C) Schematic depicting Path-GFP intensity quantification where the average GFP intensity of the glial membrane (marked in RFP) lining the surface of each brain lobe was used to reflect BBB Path-GFP expression (described in methods). (D) Quantification of the normalised (to Fed) Path-GFP intensity at the brain surface in (A and B) (n = 8, 8). (E-F) Single-section images of *elav^C155^QF>QUAS- prosRi* tumour brains with *pathRi^KK^*(F) overexpressed in PG using *GMR85G01-G4*, compared to *mCherryRi* (E) at 120hALH. Glia: Repo; NBs: Dpn. (G-H) Quantifications of the normalised (to *mCherryRi*) glial number (G) and Dpn voxels (H) of each circled brain lobe in (E-F) (n = 24, 22). (I-J) Quantifications of the normalised (to control fed condition) glial number (I) and Dpn voxels (J) of *elav^C155^QF>QUAS-prosRi* tumour brain lobes, where *path.A* was overexpressed in PG using *GMR85G01-G4*, compared with *control* (*GMR85G01- G4 X w1118*) under Fed vs -Yeast (n = 7, 12, 8, 6). Yeast dropout: 72-120hALH; Dissection: 120hALH. (K-L) Quantifications of the normalised (to Fed) glial number (K) and Dpn voxels (L) of each *elav^C155^QF>QUAS-prosRi* tumour brain lobes under Fed, dropout of Yeast (-Y) and -Yeast with 5xTrp or 5xAla or 5xPro conditions from 72-120hALH (n = 10, 14, 12, 10, 6). (M) Quantifications of the normalised (to *mCherryRi*) AA concentration in *elav^C155^QF>QUAS-prosRi* tumour brains, where *mCherryRi* or *pathRi^KK^* was overexpressed in PG using *GMR85G01-G4*. Data information: ALH = hours after larvae hatching. Brain lobes are circled with yellow dashed lines. Scale bar = 50μm. Error bar represents SEM. In (D): Mann–Whitney test, (***) P = 0.0002. In (G): Mann–Whitney test, (****) P < 0.0001. In (H): Mann–Whitney test, (****) P < 0.0001. In (I-J): Two-way ANOVA were used to analyse whether the effect of NR on glial number or tumour size is affected by *path.A* overexpression in PG (significance indicated by interaction P value). In (I): interaction P = 0.0013. In (J): interaction P = 0.0330; Mean± SEM, and statistical results including multiple comparisons are displayed in S1G-H Table. In (K): One-way ANOVA, (****) P < 0.0001, (ns) P = 0.4251, (ns) P > 0.9999, (ns) P = 0.0964. In (L): Kruskal-Wallis test, (****) P < 0.0001, (ns) P > 0.9999, (ns) P > 0.9999, (ns) P = 0.1124. In (M): His: unpaired t-test, (**) P = 0.0082; Lys: unpaired t-test, (***) P = 0.0003; Met: unpaired t-test, (**) P = 0.0049; Trp: unpaired t-test, (*) P = 0.0480; Val: unpaired t-test, (**) P = 0.0032; Leu: Welch’s t-test, (*) P = 0.0110; Thr: unpaired t-test, (*) P = 0.0266; Asn: unpaired t-test, (***) P = 0.0001; Ser: unpaired t-test, (****) P < 0.0001; Tyr: unpaired t-test, (**) P = 0.0068.

### Path is necessary and sufficient for PG expansion and tumour growth

It was previously shown that Path is required for NB proliferation [7]. To assess whether Path is required for glial expansion and tumour growth, we knocked down *path* in tumour glial cells (*repo-GAL4*) using *pathRNAi* (*KK*). This resulted in a reduction of both total glial numbers and tumour size (S7D-G Fig), and these effects were further recapitulated using a PG driver (*GMR85G01-GAL4*, Fig 4E-H), but not a CG driver (*NP2222-GAL4* [46], S7H-K Fig). Induction of *pathRNAi* (*KK*) in PG using *GMR85G01-GAL4* eliminated Path- GFP expression at the PG membrane (marked by *UAS-mRFP*, S7A-C Fig), confirming the effectiveness of the RNAi reagent. In addition, similar effects on glial number and tumour size were found using an independent *pathRNAi* line with *dcr2* driven by *repo-GAL4* (S7L-O Fig). Together, these findings suggest that Path is required in PG cells to regulate glial expansion and tumour growth.

Importantly, the overexpression of *path* (*UAS-pathA* [47]) in PG was able to partially override the effect of NR on glial and tumour growth. Compared to the control, this manipulation partially rescued the decrease in glial number and tumour size under starvation (yeast dropout) (Fig 4I-J, the number of glia and tumour size are reduced 58% and 67%, respectively, upon yeast dropout in control, and 28% and 32% in *pathA*, statistical assessment described in methods). Together, these results suggest that Path is required by the PG cells in tumour brains and its overexpression can partially enhance glial proliferation under NR. To assess whether Path is generally required in regulating gliogenesis, we knocked down *path* using *repo-GAL4* in wild-type brains. We found that *path* knockdown did not affect glial numbers in wild-type brains under fed conditions (S7P-R Fig). However, *path* knockdown reduced glial number and NB proliferation under NR (S7S-V Fig), consistent with previous findings [25]. Together, these data suggest that it is likely that glia subjected to stress (such as tumour or under NR) rely on Path, whereas wild-type glia do not.

### Path regulates the availability of amino acids in the brain

Path has been shown to have high affinity, but low capacity to transport its substrates tryptophan (Trp), proline (Pro) or alanine (Ala) [48, 49]. We found that supplementing the yeast-free food with 5xTrp, Pro or Ala from 72-120 hours ALH was insufficient to restore glial or tumour growth (Fig 4K-L), suggesting that Path is unlikely to affect tumour growth through these AAs. Next, we conducted LC-MS/MS on tumour brain samples to assess if *path* knockdown in PG can account for the reduced import of AAs in the brain under NR. In contrast to NR, where we observed a general reduction in most AAs (S4B Fig), *path* knockdown in PG did not result in reduced Trp, Pro or Ala, but instead led to a reduction in BCAAs (Leu, Ile and Val), as well as other AAs such as His, Met, Thr, Ser, Asn and Tyr (Fig 4M). Together, these data suggest that PG Path, rather than transporting its known substrates, likely regulates glial and tumour growth by controlling the transport of other AAs, including BCAAs.

### Path expression is regulated by Ilp6-InR/PI3K pathway

Next, we investigated the regulatory signals influencing Path expression. First, we evaluated the role of the PI3K pathway, a well-established mediator of cellular growth in response to nutrient availability [50]. We first assessed whether the PI3K pathway is altered in the glia of tumour brains under NR by using a FOXO-GFP reporter. Forkhead box O (FOXO) is known to translocate from the cytoplasm to the nucleus when the PI3K pathway is inhibited (Fig 5A) [51]. Under NR, we found that FOXO level was increased in the nuclei of surface glia cells in tumour brains (assessed at superficial brain sections, Fig 5B-D), suggesting that the PI3K pathway is downregulated under NR. Next, we investigated whether altering the InR/PI3K pathway influences Path expression (Fig 5A). Expression of a dominant-negative form of the Insulin Receptor (*InR^DN^*) in PG (using *GMR85GO1-G4*) significantly reduced Path-GFP levels (marked by mRFP) (Fig 5E-H). Similar results were obtained using *repo-GAL4*, and in addition to *InR^DN^*, overexpression of the PI3K adaptor protein P60 also reduced Path-GFP expression in glia (S8H Fig). Conversely, expression of a constitutively active form of *InR* (*InR^CA^*) significantly increased Path-GFP expression in PG (Fig 5E-H). Given that this manipulation delayed larval development, we further validated our result with *repo-G4*, where activation of *InR^CA^* consistently increased Path-GFP expression in glia (S8H Fig). Together, our data indicate that Path expression is positively regulated by the InR/PI3K pathway. Functionally, temporal inhibition of the PI3K pathway, achieved by overexpressing P60 in the glia of tumour brains post-CW (Fig 5I), significantly decreased glial numbers and tumour size (Fig 5J-M), mimicking the effects of NR. Additionally, temporal activation of the PI3K effector AKT (*myrAKT*) resulted in a smaller reduction in both glial number and tumour size under NR compared to the control (Fig 5N-P, the number of glia and tumour size are reduced 73% and 77%, respectively, upon NR in *mCherryRi*, and 25% and 25% in *myrAKT^OE^*, statistical assessment described in methods).

**Fig 5.**
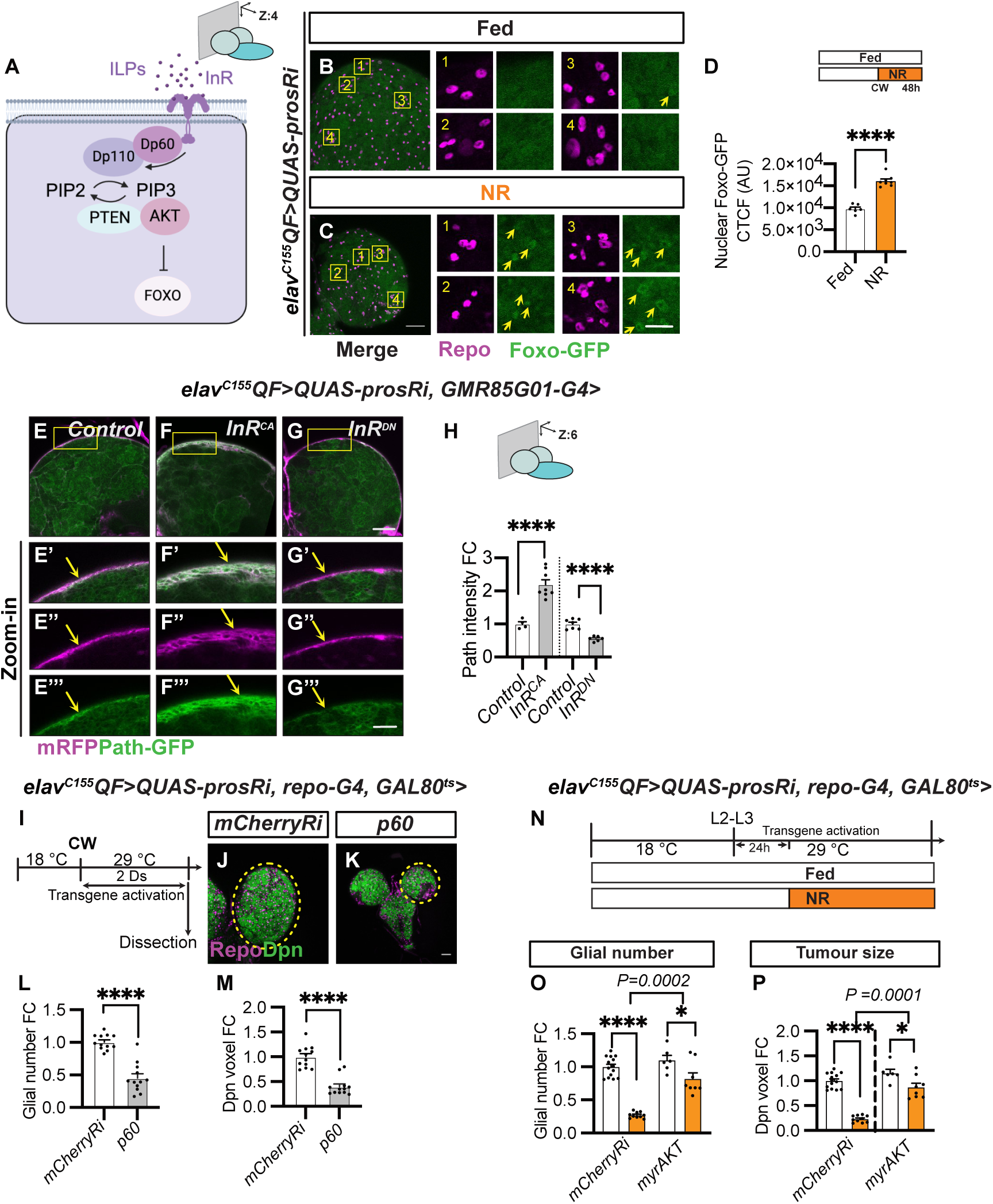
Glial Path expression is regulated by the InR/PI3K pathway. (A) Schematic depicting the PI3K pathway. The binding of Ilps to the insulin receptor (InR) activates PI3K (Dp110) and downstream signalling cascade: Dp110 first converts PIP2 to PIP3, the production of which is negatively regulated by PTEN. PIP3 then recruits AKT and allows phosphorylation of AKT at the plasma membrane, preventing the transcription factor FOXO from entering the nucleus. Created in BioRender. Dong, Q. (2025) https://BioRender.com/l19t132. (B-C) surface-section images and zoomed-in images of Foxo-GFP expression (Yellow arrows) in *elav^C155^QF>QUAS-prosRi* tumour brains with glia stained with Repo under Fed and NR. NR: 72-120hALH; Dissection: 120hALH. Scale bar = 20μm in zoomed-in images (1-4). (D) Quantification of glial nuclear Foxo-GFP intensity in (B-C, glia nucleus: Repo) (n = 6, 8). (E-G) deep-section (E-G) and zoomed-in images (E’- G’’’) of Path-GFP expression (yellow arrows) in *elav^C155^QF>QUAS-prosRi* tumour brains with *InR^DN^* or *InR^CA^* overexpressed in PG. The PG membrane is marked with *GMR85G01- G4>mRFP*. (H) Quantification of the normalised (to *control*) Path-GFP intensity at the PG membrane in (E-G) (n = 4, 8, 7, 6). The control in the first column is the same as that in S7C Fig. Scale bar = 20μm in (E’-G’’’). (I) Schematic depicting the transgene activation regime in (J-K). Tumour-bearing animals were moved from repressive 18 °C to permissive 29 °C 6-7 days after egg laying (AEL) to allow transgene activation for 2 days from CW onwards. (J-K) Single-section images of *elav^C155^QF>QUAS-prosRi* tumour brains with *p60* (K) overexpressed in glia using *repo-G4* according to the regime in (I), compared to *mCherryRi* (J). Glia: Repo; NBs: Dpn. (L-M) Quantifications of the normalised (to *mCherryRi*) glial number (L) and Dpn voxels (M) of each circled brain lobe in (J-K) (n = 12, 12). The same *mCherryRi* data was used in (L) and Fig 6Q; The same *mCherryRi* data was used in (M) and Fig 6R. (N) Schematic depicting the transgene activation and NR regime in (O-P). Genotype: *elav^C155^QF>QUAS-prosRi*, *repo-G4, GAL80^ts^* > *mCherryRi* vs *myrAKT* in (O-P). Tumour- bearing animals were moved from 18 °C to 29 °C at the L2-L3 transition to allow transgene activation only from L3 onwards. Animals were placed on either standard food or 0.42% Agar/PBS for NR 24 hours after L3 (equivalent to the timing of CW at 25°C) for 2 days before dissection. (O-P) Quantifications of the normalised (to *mCherryRi* fed condition) glial number (O) and Dpn voxels (P) of *elav^C155^QF>QUAS-prosRi* tumour brain lobes, where *myrAKT* were overexpressed in glia temporarily according to (N), compared with *mCherryRi* under Fed vs NR (n = 14, 10, 6, 8). Data information: ALH = hours after larvae hatching. Brain lobes are circled with yellow dashed lines. Scale bar = 50μm. Error bar represents SEM. In (D): unpaired t-test, (****) P < 0.0001. In (H): One-way ANOVA, (****) P < 0.0001; unpaired t-test, (****) P < 0.0001. In (L): One-way ANOVA, (****) P < 0.0001. In (M): Kruskal-Wallis test, (****) P < 0.0001. In (O-P): Two-way ANOVA were used to analyse whether the effect of NR on glial number and tumour size is affected by *myrAKT* overexpression in glia (significance indicated by interaction P value). In (O): interaction P = 0.000. In (P): interaction P = 0.0001. Mean± SEM, and statistical results including multiple comparisons are displayed in S1I-J Table.

Together, our findings suggest that NR downregulates PI3K signalling activity in the PG, which is essential for maintaining *path* expression. This regulation impacts glial number and tumour size, highlighting the role of the PI3K pathway in glia in tumour growth control.

To understand how insulin signalling is regulated by NR, we assessed the source of Insulin-like peptides (Ilps), which are known to activate insulin signalling. Insulin-producing cells (IPCs) are known to be the main source of Ilps, which regulate systemic growth [52, 53]. We previously showed that the expression of *p60* in the IPCs leads to a reduction in IPC size, consequently reducing the release of Ilps into the hemolymph [24]. Similarly, this manipulation in tumour-bearing animals resulted in smaller pupae (S8A, B and C Fig) and halted the growth of peripheral tissues (e.g. salivary glands, S8A’, B’ and D Fig). However, this unexpectedly led to an increase in glial number and tumour size, in contrast to the effects observed with NR (S8A’’, B’’, E and F Fig). Our data suggests that systemic Ilps reduction is not mediating the effect of NR on glial and tumour growth.

Other sources of Ilps include adipose tissue (the fat body) [54, 55] and glial cells within the CNS [31, 56]. We found that Ilp6 is expressed at lower levels in the fat body and glial cells of tumour-bearing animals compared to controls (examined via *Ilp6-GAL4>nGFP* [57]) (Fig 6A-J, S8G Fig). Under NR, while fat body Ilp6 levels became elevated in control animals (Fig 6A-B’ and E) [54, 57], they remained unaltered in tumour-bearing animals (Fig 6C-D’ and E). Conversely, Ilp6 expression in glia, especially in CG but not PG, was reduced upon NR in both control and tumour animals (Fig 6F-J and S8G Fig). To determine the functional importance of Ilp6 in the fat body and glial cells, we inhibited Ilp6 in these tissues post-CW (Fig 5I). Inhibiting Ilp6 in the fat body led to a significant reduction in both glial number and tumour size (Fig 6K-N). Similarly, expressing *Ilp6 RNAi* in glia post-CW mildly reduced glial number and tumour size (Fig 6O-R). These findings demonstrate that both fat body- and glial-derived Ilp6 play important roles to maintain insulin signalling in the PG to maintain its proliferation and in turn support tumour expansion.

**Fig 6.**
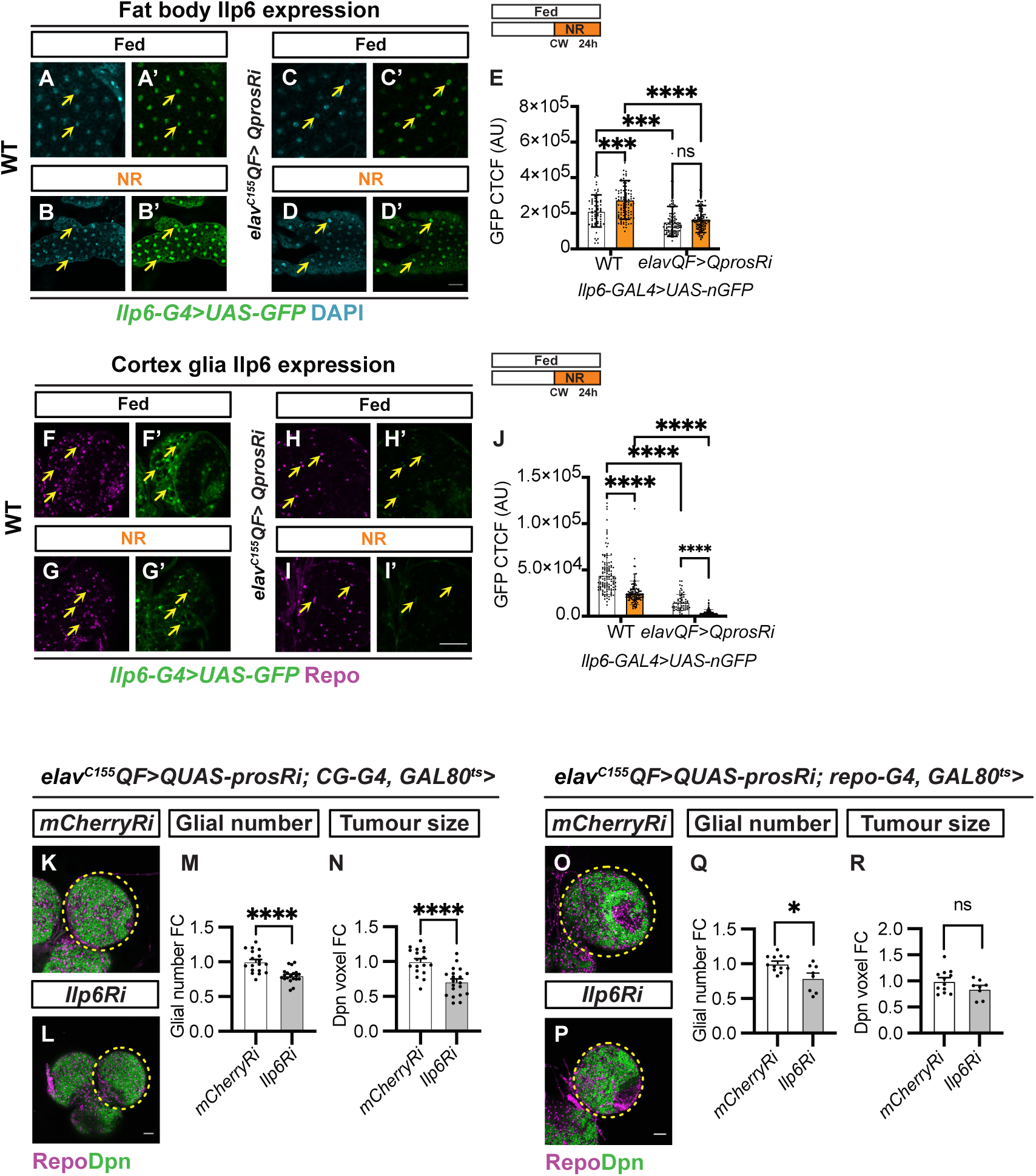
The glial PI3K pathway is sustained by Ilp6 secreted from the fat body and glia. (A-D’) Single-section images of Ilp6 expression (*Ilp6-G4>nGFP*) in fat body (FB) of wild- type and *elav^C155^QF>QUAS-prosRi* tumour brains under fed and NR conditions (yellow arrows). Wild-type and tumour-bearing animals were starved after CW (65hALH and 68hALH, respectively). The cell nucleus was marked with DAPI. (E) Quantifications of GFP intensity (CTCF) in FB nucleus in (A-D’) (n = 63, 104, 102, 99 cells from 7, 9, 9, 9 FBs) (F-I’) Single-section images of Ilp6 expression (*Ilp6-G4>nGFP*) in CG of wild-type and *elav^C155^QF>QUAS-prosRi* tumour brains under fed and NR conditions (yellow arrows). Wild-type and tumour-bearing animals were starved after CW (65hALH and 68hALH, respectively). Cortex glial cells were marked with Repo and distinguished from other glial cell types based on location. (J) Quantifications of GFP intensity (CTCF) in CG nucleus in (F-I’) (n = 120, 115, 60, 96 cells from 12, 12, 6, 10 brain lobes). (K-L) Single section images of *elav^C155^QF>QUAS-prosRi* tumour brains with *Ilp6Ri* (L) overexpressed specifically in the fat body using *CG-G4* according to the regime in (Fig 5J), compared to *mCherryRi* (K). Glia: Repo; NBs: Dpn. (M-N) Quantifications of the normalised (to *mCherryRi*) glial number (M) and Dpn voxels (N) of each circled brain lobe in (K-L) (n = 18, 20). (O-P) Single section images of *elav^C155^QF>QUAS-prosRi* tumour brains with *Ilp6Ri* (P) overexpressed in glia using *repo-G4* after CW according to the regime in (Fig 5J), compared to *mCherryRi* (O). Glia: Repo; NBs: Dpn. (Q-R) Quantifications of the normalised (to *mCherryRi*) glial number (Q) and Dpn voxels (R) of each circled brain lobe in (O-P) (n = 12, 8). Data information: ALH = after larvae hatching. Brain lobes are circled with yellow dashed lines. Scale bar = 50μm. Error bar represents SEM. In (E): Two-way ANOVA were used to analyse whether the effect of NR on FB Ilp6 expression is different in the tumour brains compared with wild-type brains (significance indicated by interaction P value). Interaction P = 0.0129. Multiple comparisons: WT fed vs NR: (***) P = 0.0001; Tumour fed vs NR: (ns) P = 0.7166; WT fed vs Tumour fed: (***) P = 0.0003; WT NR vs Tumour NR: (****) P < 0.0001.In (J): Two-way ANOVA were used to analyse whether the effect of NR on CG Ilp6 expression is different in the tumour brains compared with wild-type brains (significance indicated by interaction P value). Interaction P = 0.0070. Multiple comparisons: WT fed vs NR: (****) P < 0.0001; Tumour fed vs NR: (****) P < 0.0001; WT fed vs Tumour fed: (****) P < 0.0001; WT NR vs Tumour NR: (****) P < 0.0001. In (M): unpaired t-test, (****) P < 0.0001. In (N): unpaired t-test, (****) P < 0.0001. In (Q): One-way ANOVA, (*) P = 0.0437. In (R): Kruskal-Wallis test, (ns) P = 0.7729.

### Path regulates PG proliferation and tumour growth via mTOR/S6K pathway

Path/SLC36A4 transporter has been associated with growth regulation through the mTOR/S6K signalling pathway (Fig 7A) in *Drosophila* wings and mice retinal pigmented epithelium [49, 58, 59]. To determine whether this relationship extends to glial cells in brain tumours, we overexpressed the dominant negative forms of S6K (via *S6K^KQ^*) and mTOR (*Tor^TED^*) in tumour brains using *repo-GAL4*. We found that these manipulations resulted in reductions in glial number and tumour size (Fig 7B-G). We next assessed whether activation of mTOR/S6K signalling pathway in PG is sufficient to rescue the decrease in glial number and tumour size induced by NR. We showed that temporal activation of S6K (*S6K^CA^*) or Rag (*RagA^CA^*) in PG were sufficient to induce a smaller reduction in glial number and tumour size upon NR compared to the *mCherryRi* control (Fig 7H-J, the number of glia and tumour size are reduced 54% and 65%, respectively, upon NR in *mCherryRi*, 20% and 30% in *Rag^CA^*, and 38% and 36% in *S6K^CA^*, statistical assessment described in Methods). Finally, while the activation of S6K did not change total glial number and tumour size, it resulted in a rescue of both parameters upon *path* knockdown (Fig 7K-L). Furthermore, inhibition of mTOR (*Tor^TED^*) using *repo-GAL4* did not significantly affect Path-GFP expression in the surface glial of tumour brains (Fig 7M-O), suggesting that mTOR likely functions downstream of Path to regulate glial proliferation and tumour growth. However, we were not able to assess the effect of Path knockdown on mTOR and S6K activity due to technical limitations (the antibody against pS6K showed it has very low expression in glial cells). Collectively, our data suggest Path regulates PG proliferation potentially via the mTOR-S6K pathway, and its expression level is regulated by the fat body and glia-derived Ilp6 and the InR/PI3K pathway.

**Fig 7.**
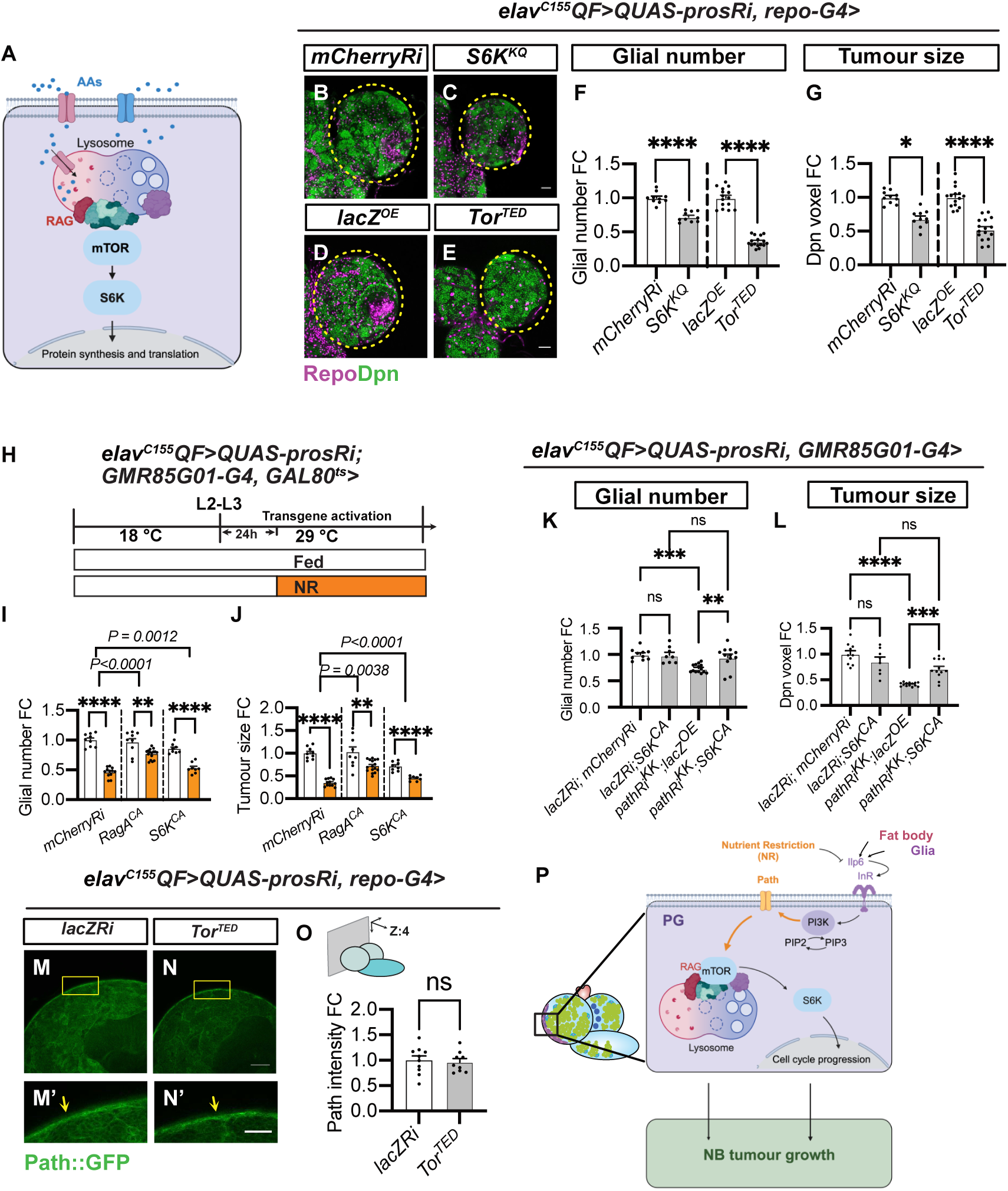
Path regulates glial proliferation via the mTOR-S6K pathway. (A) Schematic depicting the activation of the mTOR-S6K pathway by AAs through Rag GTPases. Created in BioRender. Dong, Q. (2025) https://BioRender.com/a373zo9 (B-E) single section images of *elav^C155^QF>QUAS-prosRi* tumour brains, where *mCherryRi, S6K^KQ^*, *lacZ* and *Tor^TED^* were overexpressed in glia using *repo-G4*. Glial cells are stained with Repo, and NBs with Dpn. (F-G) Quantifications of the normalised (to *mCherryRi* or *lacZ^OE^*) glial number (F) and Dpn voxels (G) of each circled brain lobe in (B-E) (n = 10, 10, 15, 16). (H) Schematic depicting the transgene activation and NR regime in (I-J). Genotype: *elav^C155^QF>QUAS-prosRi*, *GMR85G01-G4, GAL80^ts^*> *mCherryRi* vs *Rag^CA^* vs *S6K^CA^*in (I- J). Tumour-bearing animals were moved from repressive 18 °C to permissive 29 °C at the L2-L3 transition to allow transgene activation from L3 onwards. Animals were placed on either standard food or 0.42% Agar/PBS for NR 24 hours after L3 (equivalent to the timing of CW at 25°C) for 2 days before dissection. (I-J) Quantifications of the normalised (to *mCherryRi* fed condition) glial number (I) and Dpn voxels (J) of *elav^C155^QF>QUAS-prosRi* tumour brain lobes, where *Rag^CA^*or *S6K^CA^* were overexpressed in PG temporarily according to (H), compared with *mCherryRi* under Fed vs NR (n = 9, 12, 9, 16, 8, 8). (K-L) Quantification of normalised (to *lacZRi; mCherryRi*) glial number (K) and Dpn voxels (L) of each *elav^C155^QF>QUAS-prosRi* tumour brain lobe, where *lacZRi; S6K^CA^*, *pathRi^KK^*; *lacZ^OE^* and *pathRi^KK^*; *S6K^CA^* were overexpressed in PG, compared with *lacZRi; mCherryRi*. The glial number and Dpn voxels were normalised to the control (*lacZRi; mCherryRi*) (n = 10, 8, 14, 12). Glial cells: Repo; NBs: Dpn. (M-N) Single-section (M and N) and zoomed-in images (M’ and N’) of Path-GFP expression in *elav^C155^QF>QUAS-prosRi* tumour brains, where *Tor^TED^* was overexpressed in glia using *repo-G4*, compared with *mCherryRi* at the brain surface (Yellow arrows) at 120hALH. (O) Quantification of the normalised (to *mCherryRi*) Path-GFP intensity at the brain surface in (M-N) (n = 10, 9). (P) Schematic depicting the working model. Nutrient Restriction (NR) inhibits perineural glial (PG) proliferation, which consequently limits brain tumour growth. We show that NR inhibits the expression of fat body and glia-derived Ilp6, causing downregulation of the InR-PI3K pathway in PG. This results in a reduction in Path expression and the downstream mTOR-S6K pathway, causing a slowdown of PG proliferation and tumour growth. Data information: ALH = after larvae hatching. Brain lobes are circled with yellow dashed lines. Scale bar = 50μm. Error bar represents SEM. In (F): One-way ANOVA, (****) P < 0.0001; Welch’s t-test, (****) P < 0.0001. In (G): Kruskal-Wallis test, (*) P = 0.0391; unpaired t-test, (****) P < 0.0001. In (I-J): Two-way ANOVA were used to analyse whether the effect of NR on glial number or tumour size is affected by *Rag^CA^* or *S6K^CA^* overexpression in glia (significance indicated by interaction P value). In (I): interaction P < 0.0001, interaction P = 0.0012. In (J): interaction P = 0.0038, interaction P < 0.0001. Mean± SEM, and statistical results including multiple comparisons are displayed in S1K-L Table. In (K): One-way ANOVA, (ns) P = 0.9998; (***) P = 0.0002; (***) P = 0.0021; (ns) P = 0.9451. In (L): One-way ANOVA, (ns) P = 0.3106; (****) P < 0.0001; (***) P = 0.0003; (ns) P = 0.2613. In (O): Welch ANOVA test, (ns) P = 0.9320.

## Discussion

Upon the transformation of normal cells into cancer cells, metabolic rewiring occurs to meet the increased proliferative demands of the tumour [1, 60, 61]. Limiting nutrient availability in the microenvironment impacts cell growth and proliferation in both physiological and pathological contexts. However, strategies targeting the altered metabolism of tumours remain limited [62], partly because different tumours and cell types within the same tumour exhibit distinct metabolic vulnerabilities [63]. Understanding cell-type-specific metabolic dependencies within a tumour is therefore critical for developing effective therapeutic approaches.

This study leverages the QF-QUAS and GAL4-UAS binary expression systems to investigate the role of nutrients in modulating brain tumour growth, where we were able to genetically manipulate key regulators in the PG cells of the BBB (using the GAL4-UAS system) and assess the effects on the tumours (generated by the QF-QUAS system). We demonstrate that NR reduces brain tumour growth by downregulating the expression of the SLC36A4 amino acid transporter Path specifically in PGs of tumour-bearing brains (Fig 7P). Under normal conditions, Path expression in PGs of tumour brains is similar to that of its wild-type counterparts; however, NR triggers the downregulation of Path exclusively in tumour brains, leading to tumour growth inhibition. Functionally, Path is both necessary and sufficient to sustain PG proliferation, and its expression can override the inhibitory effects of NR on tumour growth. Moreover, Path expression is regulated by InR/PI3K signalling, sustained by two distinct pools of Ilp6 derived from the fat body and glial cells. In wild-type animals, the fat body upregulates *Ilp6* under NR; however, in tumour-bearing animals, this upregulation fails to occur. Previous studies have demonstrated that 20E is sufficient to induce *Ilp6* expression [54]. We have also shown that tumour-bearing animals exhibit a deficit in systemic 20E levels [27].Therefore, it is likely that the absence of 20E prevents the upregulation of *Ilp6* under NR conditions in tumour brains, a hypothesis we will test in follow-up studies.

In this study, we have identified the PG of the BBB as a critical nutrient transportation hub for modulating the response of NB tumours to NR. We have also found that the growth of these PG cells is directly controlled by the amino acid transporter Path. While Path has been shown to regulate overall body size and NB proliferation in developmental contexts, it is not required for the growth of other tissues, such as oenocytes, fat body, and salivary glands [25, 47, 49]. In tumour contexts, Path was previously shown to be upregulated to enhance the import of AAs, thereby promoting the growth of wing disc tumours [58]. Our results extend this finding, supporting a growth-promoting role for Path in the PG of tumour brains, highlighting its essential function in tumour growth regulation under nutrient-restricted conditions.

Despite previous findings suggesting that Path is essential for scavenging proline to drive tumour growth in high-sugar-dependent tumorigenesis [58], our study did not identify proline as the primary substrate for Path in this context. Unlike other SLC36 transporters, which typically function by coupling AA transport with proton exchange [64], SLC36A4/Path operates at neutral pH and exhibits a high affinity but low capacity for transporting AA substrates like Trp, Pro, and Ala [48, 49, 65]. In line with this, we show that supplementing animals with Trp, Pro or Ala was not able to increase PG number or tumour size under NR. Therefore, these AA are unlikely to be the rate-limiting substrates for Path in driving brain tumour growth. Instead, inhibition of Path in the PG of tumour brains led to a reduction in the concentration of BCAAs, suggesting that BCAAs may play a more critical role in the regulation of tumour growth, likely through mechanisms yet to be fully elucidated.

Previously, it was shown that Paneth cells act as nutrient-sensing niche cells in the mammalian gut, linking dietary cues to intestinal stem cell (ISC) expansion and tissue homeostasis [66]. For instance, the NR-induced alterations in glial cell proliferative dynamics, suggest that the tumour microenvironment, much like the intestinal niche, is highly responsive to metabolic and dietary changes. These insights highlight a promising avenue for nutrient-based interventions to modulate tumour growth by leveraging the nutrient-sensing capabilities of niche cells.

In summary, our study demonstrates that targeting the Path transporter at the BBB can effectively limit brain tumour growth, while sparing normal brain tissue. Given that the BBB presents a significant challenge for drug delivery in brain cancer treatment, our findings offer a novel strategy to restrict tumour growth without the need for drugs to cross the BBB. AA transporters may serve as potential therapeutic targets given their role in cancer cell metabolism, and their involvement in the exchange of amino acids between the tumour and stroma. Further research into the specific mechanisms and vulnerabilities of these transporters could lead to the development of novel anticancer therapies that exploit the metabolic dependencies of cancer cells.

## Materials and Methods Fly husbandry and strains

Fly strains were reared on standard food at room temperature. The standard food contains 0.42% food-grade agar, 5.25% frozen yeast, 4.38% glucose and 4.97% Polenta. Crosses for overexpression and knockdown experiments were set up at 25 °C, and after 24 hours, the progenies were moved to 29 °C, unless otherwise stated.

The fly stocks used in this study include: *repo-GAL4* (BDSC7415), *elav-QF2^C155^* (BDSC66466), *elav-GAL4^C155^* (BDSC458), *Insc-GAL4, wor-GAL4* (from Alex Gould lab), *wrapper-GAL4* (from Owen Marshall lab), *GMR85G01-GAL4* (BDSC40436), *NP2222-GAL4* (Kyoto112830), *CG-GAL4* (BDSC7011), *Ilp2-GAL4* (from Christen Mirth lab), *Ilp6-GAL4* (KDRC103877), *tubGAL80^ts^*(BDSC7108 and 7017), *path-GFP* (from Sarah Bray lab and Jay Parrish lab) [45], *dFOXO-GFP* (generated in this study), *UAS-mCherryRNAi* (BDSC35785), *QUAS-prosRNAi* [27], *UAS-prosRNAi* (BDSC42538), *UAS-bratRNAi* (BDSC34646), *UAS- LeuRSRNAi* (BDSC34483), *UAS-pathRNAi* (VDRC100519 and BDSC64029), *UAS- Ilp6RNAi* (BDSC33684), *UAS-myrRFP* (BDSC), *UAS-dcr2*, *UAS-lacZ, UAS-htl^ACT^* (BDSC5367), *UAS-aPKC^CAAX^* (Helena Richardson Lab), UAS*-Cdk4, UAS-CycD; +/TM6B* [67], *UAS-S6K^KQ^* (BDSC6911), *UAS-Tor^TED^* [68], *UAS-path.A* (from Susumu Hirabayashi lab and Jay Parrish lab) [47], *UAS-RagA^CA^* (from Thomas Neufeld lab), *UAS-S6k^CA^* (BDSC6914), *UAS-InR^DN^*(BDSC8252), *UAS-InR^CA^* (BDSC8263), *UAS-p60* [24], *UAS-myrAKT* [69], *ey- FLP1;QUAS-scribRNAi;QUAS-Ras^V12^/CyOQS; act>CD2>QF2, UAS-RFP/TM6QS* [70], *UAS-GFP, UASp-CFP.E2f1.1-230, UASp-Venus.NLS.CycB.1-266/TM6B* (FUCCI, BDSC55114).

## Generation of dFOXO-GFP

Foxo IO@CTV-PaxCherry: vector was used to tag dFoxo with eGFP in the C-terminus. To generate the dFoxo-GFP fusion, we followed the protocol described in [71]

Using CRISPR/Cas 9, we inserted eGFP just before the stop codon in exon 10 (foxo RB). Primers and vectors sequences used for the design of IO@CTV-Pax-Cherry are available upon request.

## Dietary manipulations

For NR experiments, larvae were raised on our standard food at 25°C and selected at the L2- L3 moulting stage within a 3-hour time window based on morphological features, unless otherwise stated. Larvae were transferred to either standard food or NR, -Yeast, - Carbohydrate, +2xYeast, CDD, CDD-Leu, CDD-Ile and CDD-Met diet (see S1M Table) in a *Drosophila* vial from either 48 or 72hALH.

## RNA sequencing

Total RNA from 5 dissected larval brains at 120hALH was pooled and extracted using Direct-zol RNA Microprep Kits (ZYMO Research, #R2061) according to the manufacturer’s instructions. 5 biological replicates were prepared for each condition (Fed and NR, NR: 72- 120hALH). RNA quantity and quality were assessed using Qubit and Agilent TapeStation.

Two samples from NR were excluded due to low concentration or low quality. cDNA Library was prepared using the QuantSeq 3’ mRNA-Seq Library Prep Kit (Lexogen). The generated reads were aligned to the *Drosophila melanogaster* genome assembly Release 6 (Dm6) and were checked with FastQC (usegalaxy.org). Mapped reads were converted to counts using featureCounts (usegalaxy.org) and loaded to Degust (https://degust.erc.monash.edu/) for MDS plots and differential gene expression analysis using limma-voom. Differentially expressed genes were matched to genes annotated as transporters in GLAD (https://www.flyrnai.org/tools/glad/web/) [72]. Genes that are significantly downregulated (FDR≤ 0.05, FC ≥ 1.5) were uploaded to FlyEnrichr [73, 74] for KEGG pathway enrichment analysis. The volcano plot was generated using GraphPad Prism 9.0.

## Quantitative reverse transcription PCR

Total RNA from 5 dissected wild-type larval brains was pooled and extracted using Direct-zol RNA Microprep Kits (ZYMO Research, #R2061), followed by reverse transcription into cDNA using ProtoScript II First Strand cDNA Synthesis Kit (NEB, #E6560S) according to the manufacturer’s instructions. 3 biological replicates were prepared for each condition (Fed and NR, NR: 63.5-96hALH). The qPCR was performed using Fast SYBR Green master mix reagent (Applied Biosystems, #4385612), on the stepOnePlus real-time PCR system (Applied Biosystems). mRNA abundance was normalised to rpl32 and calculated using the 2^-ΔΔCt^ method. Primers for *rpl32*[75]: Forward: CCGCTTCAAGGGACAGTATCTG; Reverse: ATCTCGCCGCAGTAAACGC. Primers for *LeuRS* (primer pair: PP36944, FlyPrimerBank, https://www.flyrnai.org/flyprimerbank): Forward: TGGCAAATTATGCAGAGTCTCG; Reverse: TGGGAAGTAGTTAAGCCAGTGTT.

## Metabolite measurements

For hemolymph sample collection, hemolymph from five tumour-bearing larvae (at 120hALH) was pooled and released into 115μl of ultrapure water before snap-freezing in liquid nitrogen. Five biological replicates were collected for each condition (Fed and NR, NR:72-120hALH). The metabolites were extracted using 100% methanol with 20μM of the internal standards (methionine sulfone and 2-morpholinoethanesulfonic acid), chloroform and acetonitrile [76]. Metabolites were quantified by ultra-performance liquid chromatography– tandem mass spectrometry (LCMS-8060NX, Shimadzu. The metabolite concentrations were normalised to 2-morpholinoethanesulfonic acid, and the protein amount in the hemolymph.

One sample from (NR condition) was excluded from the analysis due to variance in the internal standard detection. For brain sample collections, six brains were dissected in cold PBS and pooled into 115μl of ultrapure water in a 1.5ml Eppendorf tube before a brief homogenisation (2 x10 seconds) on ice. The tube was then snap-frozen in liquid nitrogen and stored at -80°C before extraction and detection as described above. The concentrations of metabolites per brain were plotted in S4B Fig and Fig 4M.

## Immunostaining

Larval brains and salivary gland (SG) were dissected in PBS, fixed for 20 minutes in 4% formaldehyde (Sigma-Aldrich, #F8775) in PBS and washed in 0.5% PBST (PBS + 0.5% TritonX-100 (Sigma-Aldrich, #T8787), 3X 10 minutes). The larval fat body was fixed for 45 minutes and washed with 0.2% PBST. The tissues were stained with primary and secondary antibodies overnight at 4 °C. Samples were mounted in 80% glycerol in PBS or Prolong^TM^ Diamond Antifade Mountant (Invitrogen, #P36961) for image acquisition. The primary antibodies used were mouse anti-Repo (1:50; DSHB, 8D12), mouse anti-Mira (1:50; gift of Alex Gould), rat anti-Mira (1:100, Abcam, #ab197788), rat anti-Dpn (1:200; Abcam, 195172), rabbit anti-pH3 (1:200, Cell Signaling, 3377), mouse anti-Elav (1:50; DSHB, 9F8A9), rabbit anti-Dcp-1-cleaved (1:100, Cell Signaling, 9578S). Secondary donkey antibodies conjugated to Alexa 555, 488 and Alexa 647 (Molecular Probes) were used at 1:500. Highly cross-adsorbed secondary antibodies were used to minimise cross-reactivity between mouse anti-Repo and rat anti-Dpn. DAPI and Phalloidin (both from Molecular Probes) were used at 1:1,000. Images were collected on an Olympus FV3000 confocal microscope.

## EdU labelling and analysis

EdU in vitro labelling was used to trace actively dividing tumour NBs in (S2D-E’ Fig). Larval brains were dissected in PBS at 120hALH and incubated in tubes with 10 μM EdU/PBS for 15 mins before fixation, and primary and secondary antibody staining (for Repo and Dpn detection). EdU detection was performed using Click-iT™ EdU Cell Proliferation Kit for Imaging, Alexa Fluor™ 647 dye (Invitrogen, #C10340) following the manufacturer’s instruction.

## Quantification and Analysis Glial cell number measurements

Glial numbers were automatically counted from the 3D reconstruction of confocal Z stacks (2-μm step-size) of each larval brain lobe using a Fiji macro “DeadEasy larval Glia” [77], followed by a FIJI plugin called 3D Objects Counter.

## Size measurements

The total volume (voxels) of the tumour cell marker was measured from the 3D reconstruction of confocal Z stacks (2-μm step-size) of each sample using Volocity software to represent the tumour size. The area of the tumour brains and salivary glands was measured from maximum projection images, and that of each fat body cell was measured from single-section images in S1C-E Fig. The pupa images in (S8A and B Fig) were collected with a brightfield microscope and the area of the pupa was measured to represent pupa size.

## Assessment of the effect of genetic manipulations on tissue sensitivity to NR

To assess whether certain genetic manipulations can alter the effect of NR on glial and tumour growth under NR, we quantified the glial number and tumour size of each genotype under Fed and NR conditions. The two-way ANOVA tests were used to understand whether the differences between NR and Fed are consistent between genotypes. A statistically significant interaction (interaction P value) means the genetic manipulation induces a significant change in the effect of NR on glial number or tumour size.

## Intensity measurements

For measurements of Path-GFP at the BBB, 8-10 lines were drawn across the brain surface glial membrane (labelled with RFP) from a single-section confocal image. Pixel values of Path-GFP and glial membrane marker RFP were generated along each line using Fiji’s “Plot Profile” tool (Fig 4C). 8-10 Path-GFP pixel value of glial membrane (reflected by the peak value of RFP pixel along the line) was recorded, and used to calculate the average of Path- GFP pixel value in glial membrane of each brain lobe (Fig 4C). For Path-GFP intensity measurements in S6J-L Fig, a single section and sum projection confocal image were used, respectively. The intensity of the signal was measured in Fiji using the following formula: CTCF (corrected total cell fluorescence) = Integrated density – (Area of the tumour x mean fluorescence of background readings) [78]. This formula was also used to calculate PG nuclear FOXO-GFP expression, and glial and fat body nuclear *Ilp6-G4>GFP* expression.

## FUCCI cell cycle analysis

FUCCI [34] was overexpressed specifically in PG using *GMR85GO1-G4* to assess the distribution of PG cells within the cell cycle (G1: marked by ECFP; S: marked by Venus; G2/M: marked by ECFP and Venus) over time in wild-type and tumour-bearing brains. The total number of PGs, as well as those in each cell cycle phase, was automatically measured using a FIJI macro we previously developed [27]. The chi-square and the two-way ANOVA tests were used to assess the difference in the distribution of cells to each cell cycle phase.

The two-way ANOVA multiple comparisons were used to assess changes in each cell cycle phase.

## pH3 index

**the NB cell cycle progression was assessed using the pH3 index, represented as** the percentage of NBs in the M phase. In Fig 1F and S7V Fig, the number of pH3^+^ Type I NBs and the total number of Type I NBs in the CB were counted manually in Fiji based on pH3 and Dpn or Mira staining. Type I NBs are distinguished from Type II NBs according to location and morphological features as previously described [79]. The % of pH3^+^ Type I NBs = the number of pH3^+^ Type I NBs / the total number of Type I NBs x 100. In Fig 3S, the ratio of NB tumour cells in M phase was assessed using the formula: The ratio of pH3^+^ NBs = the total volume of pH3 in Dpn^+^ cells / the total volume of Dpn in each brain lobe.

## EdU index

The cell cycle progression of tumour NBs was assessed using the EdU index, represented as the percentage of NBs that can incorporate EdU *ex vivo* within a 15-minute time window.

The % of EdU^+^ NBs = the total volume (voxels) of EdU in Dpn^+^ cells/ the total volume of Dpn in each brain lobe x100.

## The neuron ratio

The neuron ratio in each brain lobe was used to represent the dedifferentiation rate in S2G-I Fig). The % of Elav^+^ cells = the total volume of Elav/ the total volume of DAPI in each brain lobe x 100.

## The ratio of cells undergoing apoptosis

The ratio of cells undergoing apoptosis was assessed based on Dcp-1 staining, using the formula: The % of Dcp-1^+^ NBs = the total volume of Dcp-1 in Dpn^+^ cells / the total volume of Dpn in each brain lobe x100. The % of Dcp-1^+^ glia = the total volume of Dcp-1 in Repo^+^ cells / the total volume of Repo in each brain lobe x100

## Statistical analysis

Shapiro-Wilk test was used to assess data normality. Two-tailed unpaired student’s t-tests and One-way ANOVA tests were used to assess the difference between two groups and three or more independent groups, respectively. The non-parametric Mann-Whitney U test and Kruskal-Wallis test were used for not normally distributed data. The Welch’s correction and Brown-Forsythe correction were used for the data of unequal variance. The chi-square test was used to test whether two categorical variables are related to each other. The two-way ANOVA tests were used to understand the interaction between genotypes and nutritional status in regulating glial number and tumour size, as well as the effect of developmental timing on the percentage of PGs in each cell cycle phase.

## Acknowledgements

We are grateful to Sarah Bray, Jay Parrish, Owen Marshall, Christen Mirth, Helena Richardson, Susumu Hirabayashi, Christian F. Lehner, Thomas Neufeld, Alex Gould and Jean-Paul Vincent for the generous sharing of antibodies and fly stocks. We would like to thank Rina Okada from the Obata lab for metabolomics expertise, Khanh Nguyen, Andrew Cox and Owen Marshall for critical reading of the manuscript. We also thank the Bloomington Drosophila Stock Centre (BDSC), Vienna Drosophila Resource Centre (VDRC), Kyoto Drosophila Stock Centre (KDRC) and Developmental Studies Hybridoma Bank for fly stocks and antibodies. We would also like to thank OZDros for *Drosophila* quarantine, Peter MacCallum Center for Advanced Histology and Microscopy for microscopy assistance, and the molecular genomics core for sequencing support. QD is supported by the Peter MacCallum Cancer Foundation Grant. FO’s lab is supported by Japan society for the promotion of science (JPJSBP120249944), LYC’s laboratory is supported by funding from the NHMRC Ideas Grant (APP2011289).

## Author contributions

**Qian Dong:** Conceptualization; Formal analysis; Investigation; Methodology; Validation; Project administration; Writing-original draft; Writing-review and editing. **Edel Alvarez-Ochoa:** Conceptualization; Formal analysis; Investigation; Methodology; Validation. **Hina Kosakamoto:** Formal analysis; Methodology; Validation. **Fumiaki Obata:** Resources; Supervision. **Cyrille Alexandre:** Methodology. **Louise Y Cheng:** Conceptualization; Resources; Formal analysis; Supervision; Funding acquisition; Investigation; Methodology; Validation; Project administration; Writing-original draft; Writing-review and editing.

## Supplemental figures, figure legends and tables

**S1 Fig.**
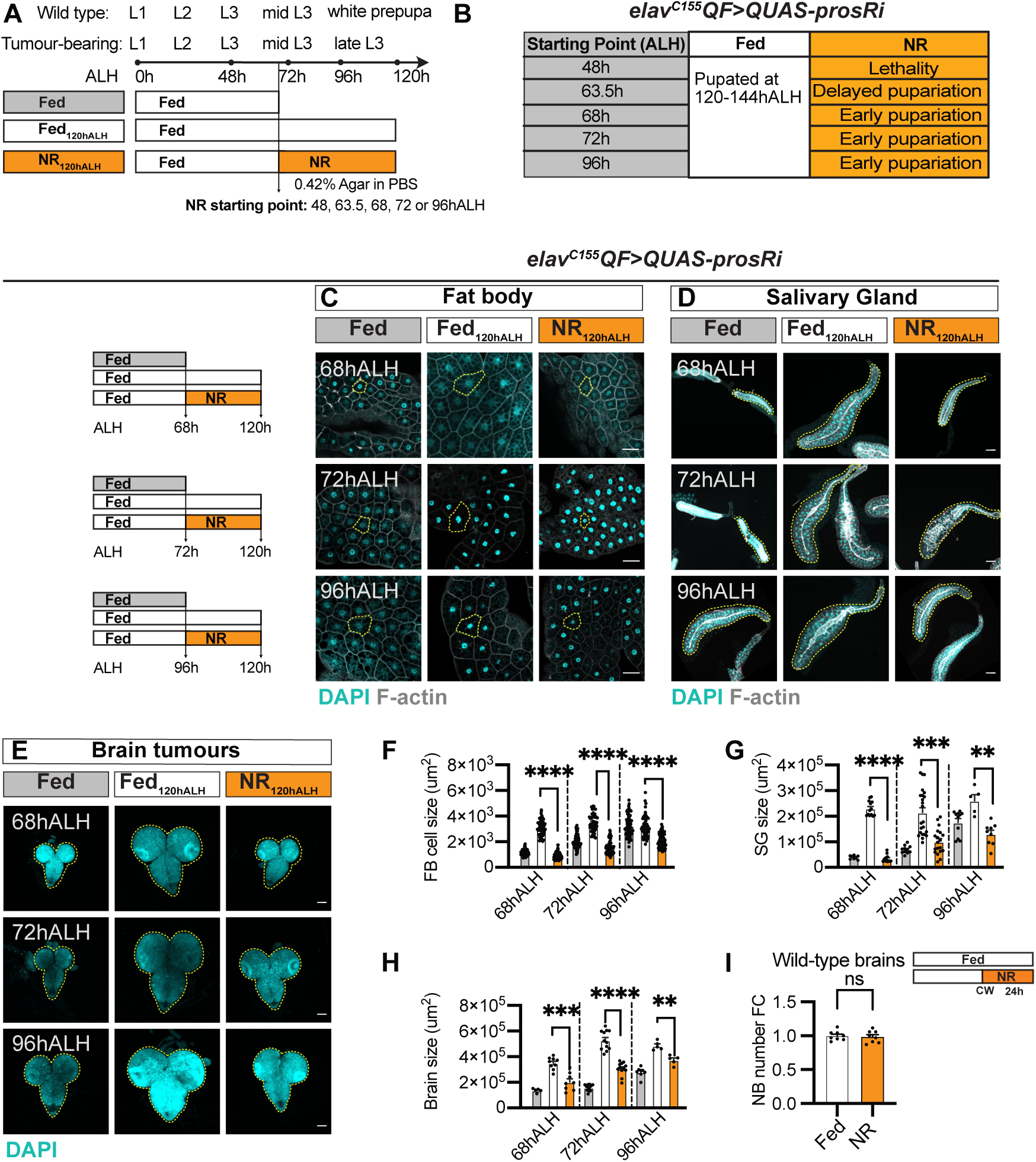
Characterisation of the effect of nutrient restriction (NR) on the growth of peripheral tissues and neuroblast (NB) tumours (related to Fig 1). (A) Schematic representation of the feeding/NR regime. Tumour-bearing larvae were fed till a certain time point before being subjected to starvation and dissected at 120hALH (NR_120hALH_) compared with larvae fed till 120hALH (Fed_120hALH_). Wild-type animals pupariated at around 96hALH while tumour-bearing animals remained as larvae until 144hALH due to developmental delay. (B) Table depicting the consequence of NR starting from 48, 63.5, 68, 72 and 96hALH on pupariation of *elav^C155^QF>QprosRNAi* tumour-bearing animals. (C-E) NR from 68, 72 and 96hALH significantly reduced the size of fat body cells, salivary glands and brain tumours (circled by yellow dashed lines). (C-D): single-section images of fat body and maximum projection images of salivary glands, marked by DAPI and Phalloidin at NR starting time point, Fed_120hALH_ and NR_120hALH_. (E): maximum projection images of brain tumours, marked by DAPI at NR starting time point, Fed_120hALH_ and NR_120hALH_. Scale bar = 100μm for (D) and (E). (F-H) Quantifications of the area of a single fat body cell circled in (C), the salivary gland circled in (D) and the whole brain circled in (E). In (F): n_68hALH_ = 105, 60, 75; n_72hALH_ = 79, 52, 68; n_96hALH_ = 74, 62, 58. In (G): n_68hALH_ = 8, 13, 12; n_72hALH_ = 11, 23, 19; n_96hALH_ = 11, 5, 9. In (H): n_68hALH_ = 5, 9, 8; n_72hALH_ = 13, 12, 12; n_96hALH_ = 8, 5, 5. Genotype: *elav^C155^QF>QprosRNAi*. (I) Quantification of type I NB number (based on Mira staining) in the CB shown in Fig. 1D-E (n = 8, 8). Data information: ALH = after larvae hatching. Scale bar = 50μm unless otherwise stated. The error bar represents SEM. In (F): Kruskal-Wallis H Test, (****) P < 0.0001; Kruskal- Wallis H Test, (****) P < 0.0001; Kruskal-Wallis H Test, (****) P < 0.0001. In (G): Kruskal-Wallis H Test, (****) P < 0.0001; Kruskal-Wallis H Test, (***) P = 0.0002; Kruskal-Wallis H Test, (**) P = 0.0011. In (H): (***) P = 0.0006; Welch’s ANOVA, (****) P < 0.0001; Welch’s ANOVA, (**) P = 0.0020. In (I): unpaired t-test, (ns) P = 0.7211.

**S2 Fig.**
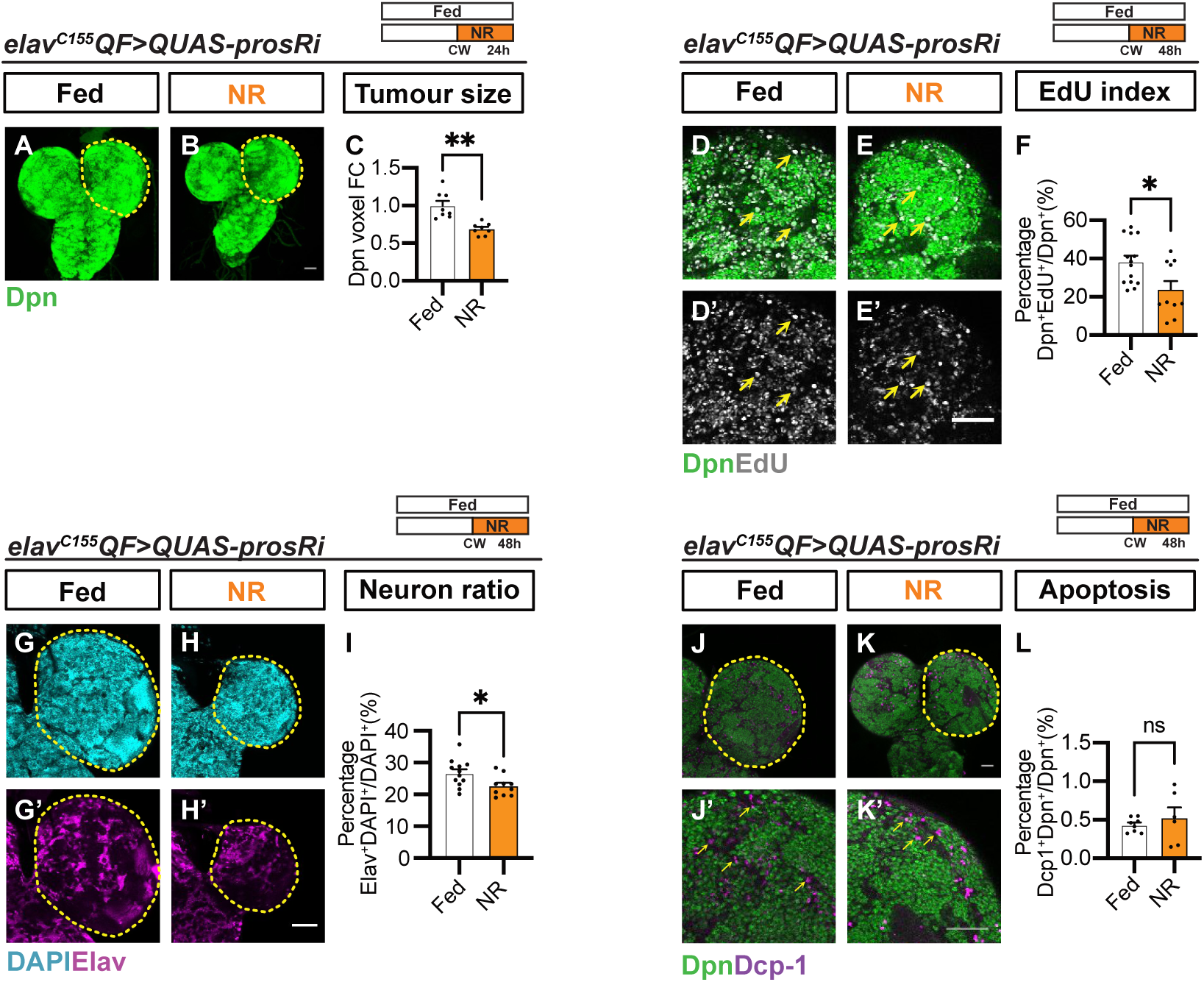
The effects of nutrient restriction (NR) on the proliferation, differentiation and apoptosis of Pros-loss-of-function neuroblast (NB) tumours (related to Fig 1). (A-B) Maximum projection images of *elav^C155^QF>QUAS-prosRi* tumour brains stained with the NB marker Dpn under Fed and NR conditions (n= 8,8). NR: 72-96hALH; Dissection: 96hALH. (C) Quantification of normalised (to Fed) Dpn voxels of circled brain lobes in (A- B). (D-E’) Zoomed-in single-section images of *elav^C155^QF>QUAS-prosRi* tumour brains labelled with EdU (yellow arrows) and Dpn under Fed and NR conditions (the same dataset from Fig. 1G-H). D and E: Dpn and EdU; D’ and E’: EdU. (F) Quantification of the percentage of NB tumours (Dpn^+^, green) that are EdU^+^ in each brain lobe of (D-E) (n = 14, 10). (G-H’) Single section images of *elav^C155^QF>QUAS-prosRi* tumour brains stained with DAPI and the neuronal marker Elav under Fed and NR conditions. (I) Quantification of the percentage of neurons among all brain cells in each brain lobe of (G-H’) (n = 12, 10). (J-K’) Single section images of *elav^C155^QF>QUAS-prosRi* tumour brains stained with the NB marker Dpn and the cell death marker Dcp-1 (yellow arrows) under Fed and NR conditions. (J’) and (K’) are zoomed-in images of (J) and (K). Dcp-1 mostly did not colocalise with Dpn in both conditions. (L) Quantification of the percentage of NBs (Dpn^+^) undergoing apoptosis (Dcp-1^+^) of each circled brain lobe in (J-K) (n = 8, 6). Data information: ALH = after larvae hatching. NR: 72-120hALH; Dissection: 120hALH unless otherwise stated. Brain lobes are circled with yellow dashed lines. Scale bar = 50μm. Error bar represents SEM. In (C): Welch’s t-test, (**) P = 0.0012. In (F): unpaired t-test, (*) P = 0.0128. In (I): unpaired t-test, (*) P = 0.0247. In (L): Welch’s t-test, (ns) P = 0.5374.

**S3 Fig.**
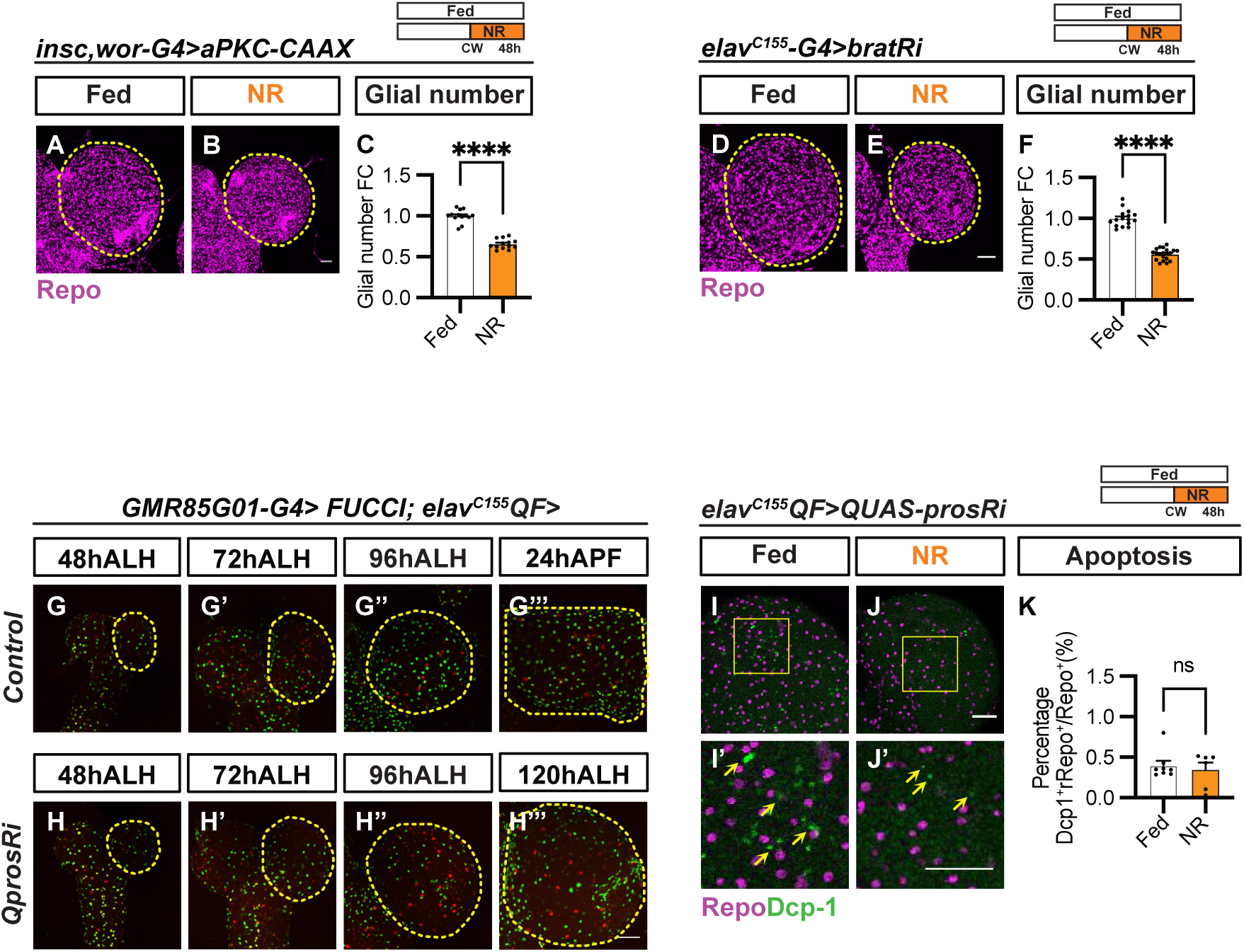
The effect of nutrient restriction (NR) on glial cell death in *elav^C155^QF>QUAS- prosRi* tumour, and the growth of other brain tumours (related to Fig 2). (A-B) Maximum projection images of *insc-G4, wor-G4>aPKC-CAAX* tumour brains stained with the glial marker Repo under Fed and NR conditions. (C) Quantification of normalised (to Fed) glial number of circled brain lobes in (A-B) (n = 13, 13). (A-C) are from the same experiment as Fig. 1J-M. (D-E) Maximum projection images of *elav^C155^-G4>bratRi* tumour brains stained with the glial marker Repo under Fed and NR conditions. (F) Quantification of normalised (to Fed) glial number of circled brain lobes in (D-E) (n = 16, 20). (D-F) are from the same experiment as Fig. 1N-Q. (G-H) Time-course single-section images of wild-type (G-G’’’) and *elav^C155^QF>QUAS-prosRi* tumour brains (H-H’’’) with FUCCI overexpressed in PG using *GMR85G01-G4* (G1-phase cells: red; S-phase cells: green; and G2/M-phase cells: yellow in Fig 2I). Time points: 48, 72, 96hALH and 24hAPF in wild-type larvae; and 48, 72, 96 and 120hALH in tumour-bearing larvae (pupate between 120-144hALH). (I-J’) Single section images of *elav^C155^QF>QUAS-prosRi* tumour brains stained with the glial marker Repo and the cell death marker Dcp-1 (yellow arrows) under Fed and NR conditions. (I’) and (J’) are zoomed-in images of I and J. Dcp-1 mostly did not colocalise with Repo in both conditions. (K) Quantification of the percentage of glial cells (Repo^+^) undergoing cell death (Dcp-1^+^ Repo^+^) among all glia in each brain lobe in (G-H) (n = 8, 6). (I-K) are from the same experiment as S2J-K Fig. Data information: ALH = after larvae hatching. APF = after pupa formation. NR: 72- 120hALH; Dissection: 120hALH unless otherwise stated. Brain lobes are circled with yellow dashed lines. Scale bar = 50μm. Error bar represents SEM. In (C): unpaired t-test, (****) P < 0.0001. In (F): unpaired t-test, (****) P < 0.0001. In (K): Mann–Whitney test, (ns) P = 0.6620.

**S4 Fig.**
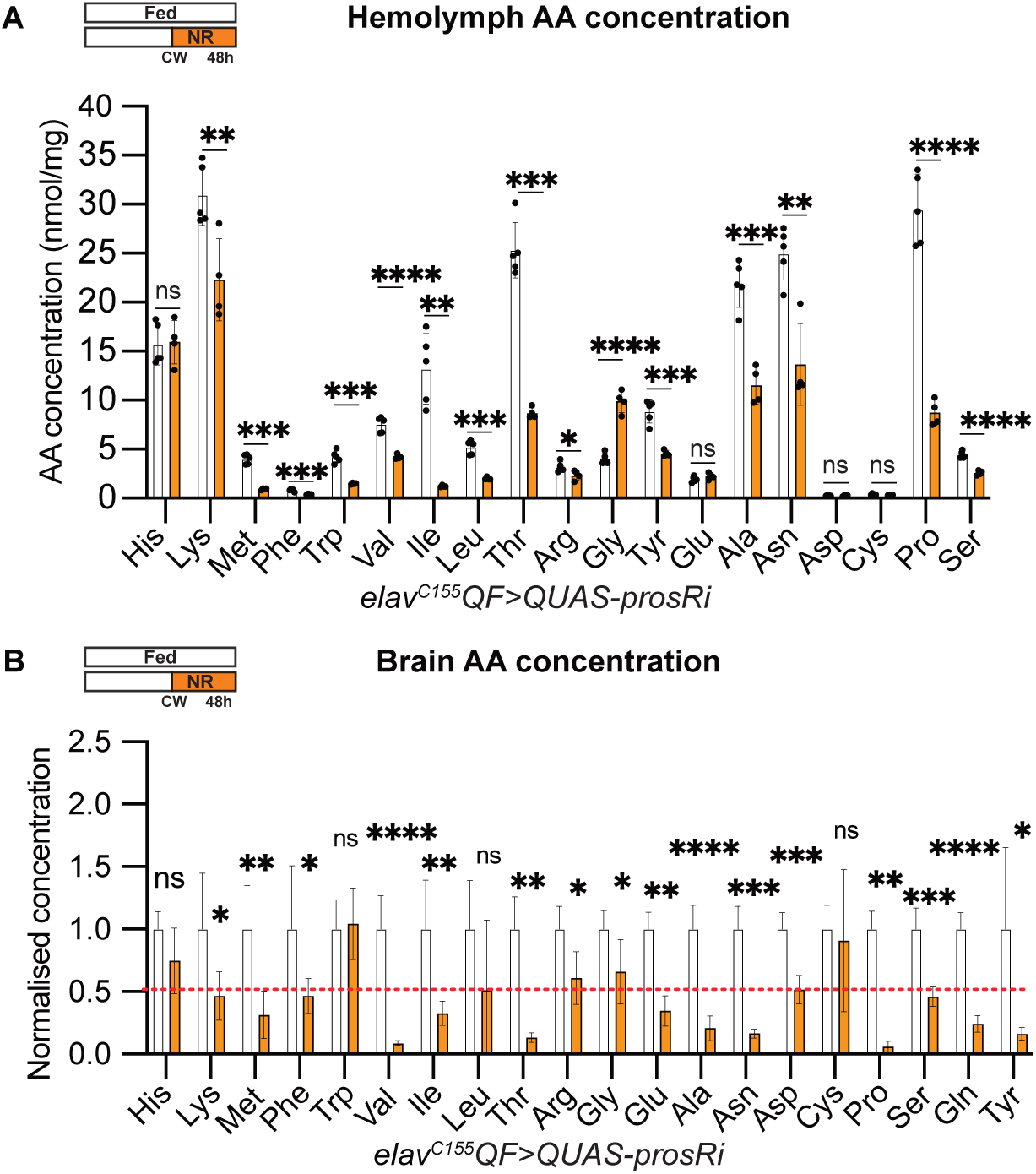
The effect of nutrient restriction (NR) on hemolymph and tumour brain amino acid content (related to Fig 3). (A) Quantification of hemolymph AA concentrations of *elav^C155^QF>QUAS-prosRi* tumour- bearing animals. NR reduced the concentration of the majority of hemolymph AAs, except for His, Glu, Asp and Cys (n = 5, 4). (B) Quantifications of the normalised (to Fed) brain AA concentration in *elav^C155^QF>QUAS-prosRi* tumour-bearing animals under Fed and NR (n= 5, 5). Data information: NR: 72-120hALH; Dissection: 120hALH unless otherwise stated. Error bar represents SEM. In (A): unpaired t-test, (ns) P = 0.8631; (**) P = 0.0091; (***) P = 0.0001; (***) P = 0.0002; (***) P = 0.0005; (****) P < 0.0001; (**) P = 0.0018; (***) P = 0.0004; (***) P = 0.0001; (*) P = 0.0322; (****) P < 0.0001; (***) P = 0.0002; (ns) P = 0.1710; (***) P = 0.0002; (**) P = 0.0016; (ns) P = 0.9419; (ns) P = 0.2580; (****) P < 0.0001; (****) P < 0.0001. In (B): Lys: unpaired t-test, (*) P = 0.0402; Met: unpaired t-test, (**) P = 0.0048; Phe: Mann–Whitney test, (*) P = 0.0317; Val: unpaired t-test, (****) P < 0.0001; Ile: unpaired t-test, (**) P = 0.0059; Thr: Welch’s t-test, (**) P = 0.0015; Arg: unpaired t-test, (*) P = 0.0138; Gly: unpaired t-test, (*) P = 0.0334; Glu: Mann–Whitney test, (**) P = 0.0079; Ala: unpaired t-test, (****) P < 0.0001; Asn: unpaired t-test, (***) P = 0.0001; Asp: unpaired t-test, (***) P = 0.0003; Pro: Mann–Whitney test, (**) P = 0.0079; Ser: unpaired t-test, (***) P = 0.0002; Gln: unpaired t-test, (****) P < 0.0001; Tyr: Welch’s t-test, (*) P = 0.0453.

**S5 Fig.**
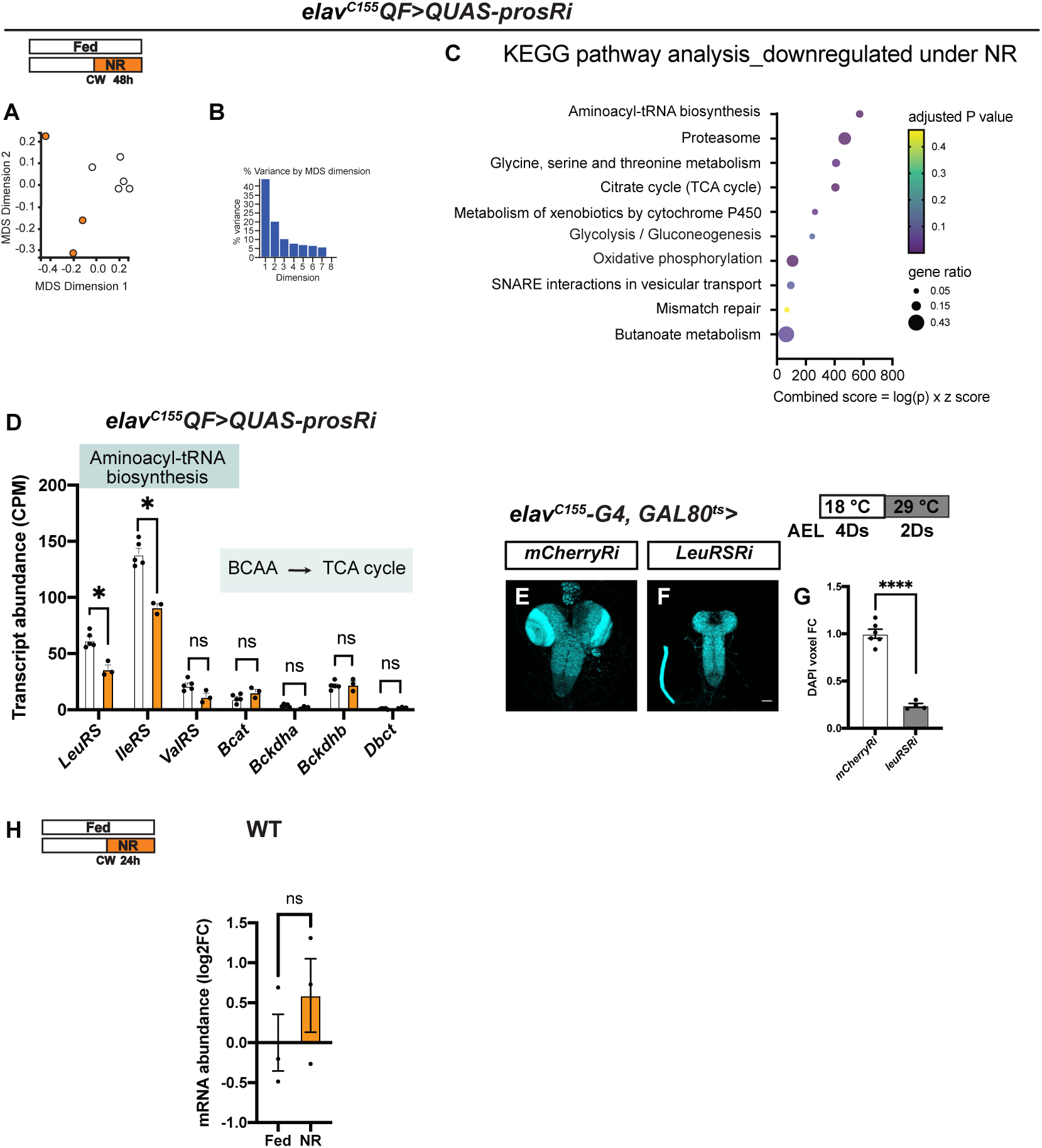
Nutrient restriction (NR) downregulates BCAA tRNA synthetases specifically in tumour brains, but not in wild-type brains (related to Fig 3). (A): Multidimensional Scaling (MDS) plot, showing gene expression profiles of *elav^C155^QF>QUAS-prosRi* brain tumour samples are separated by nutrient conditions (fed vs NR). NR: 72-120hALH. (B): The variance table displaying how much each dimension in the MDS plot accounts for the total variance in the data. (C) KEGG pathway analysis showing top 10 downregulated pathways, ranked by the FlyEnrichr combined score. (D) Plot of transcript abundance alteration of *LeuRS*, *IleRS*, *ValRS*, *Bcat*, *Bckdha*, *Bckdhb* and *Dbct* in the tumour brains under NR vs fed (from RNA seq data, n = 5,3). (E-F) Maximum projection images of wild-type brains, where *mCherryRi* or *LeuRSRi* was overexpressed in NBs using *elav^C155-^G4, GAL80^ts^.* Larvae were placed at 18°C for 4 days before being moved to 29°C for transgene activation (2 days). Brains are stained with DAPI. (G) Quantifications of the normalised (to *mCherryRi*) DAPI voxels of each brain in (E and F) (n = 6,4). (H) Quantification of mRNA level of *LeuRS* in wild-type brains under NR vs fed (by RT-qPCR, n= 3,3). NR: 63.5-96hALH. Data information: NR: 72- 120 hALH; Dissection: 120hALH unless otherwise stated. Error bar represents SEM. In (D): (*) FDR = 0.024; (*) FDR = 0.043; (ns) FDR> 0.05. In (G): unpaired t-test, (****) P < 0.0001. In (H): unpaired t-test, (ns) P = 0.3666.

**S6 Fig.**
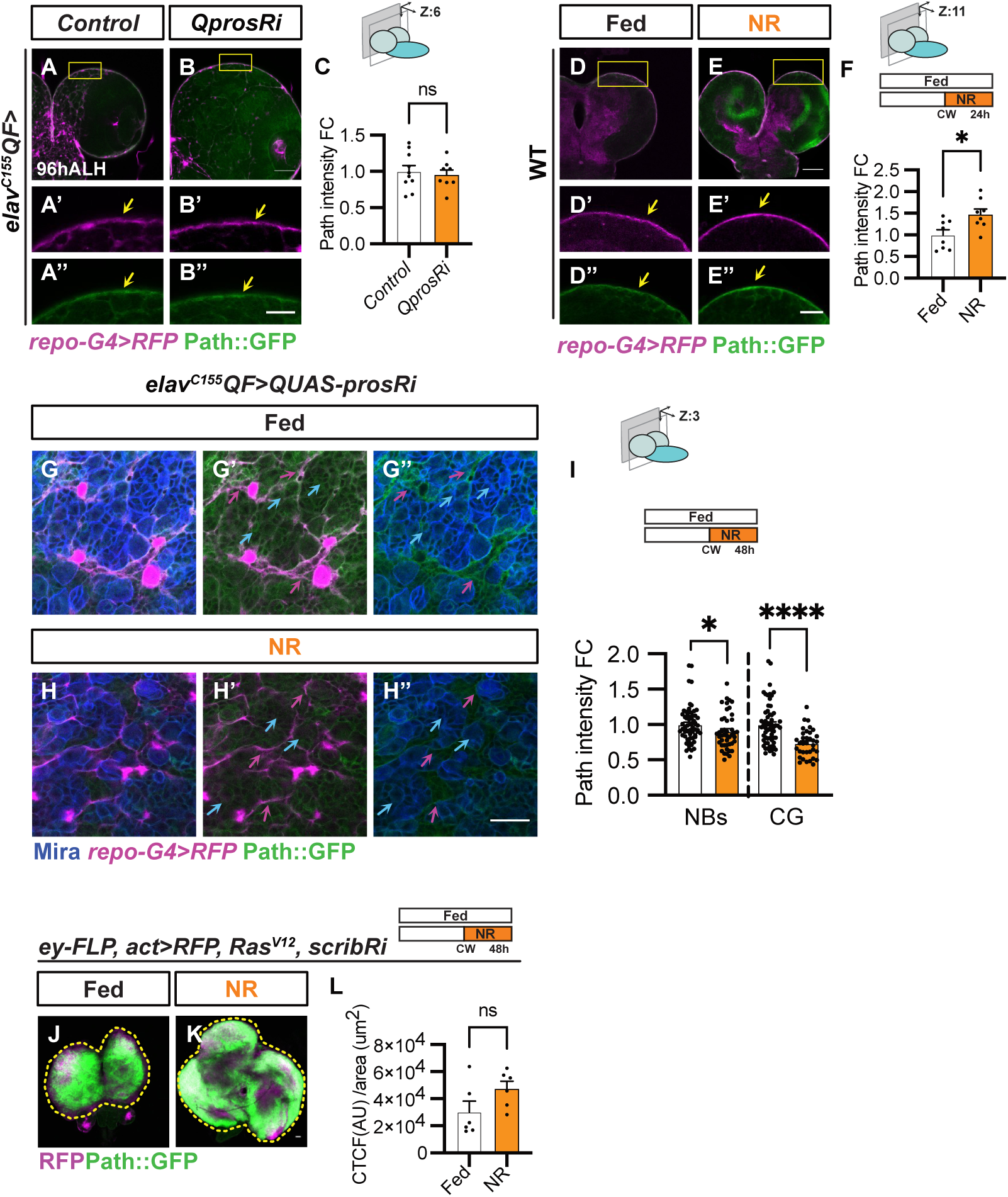
Characterisation of the effect of nutrient restriction (NR) on Path expression in other brain cell types and other tumours (related to Fig 4). (A-B) Single-section (A and B) and zoomed-in images (A’-B’’) of Path-GFP expression in control vs *elav^C155^QF>QUAS-prosRi* tumour brains at 96hALH. Glia at the brain surface are marked with *repo-G4>mRFP* and distinguished from the other glial types based on position (yellow arrows). Scale bar = 20μm in (A’-B’’) (C) Quantification of the normalised (to control) Path-GFP intensity at the BBB in (A’ and B’) (n = 10, 8). (D-E) Single-section (D and E) and zoomed-in images (D’-E’’) of Path-GFP expression in wild-type brains under Fed and NR. Glial membrane at the brain surface is marked with *repo-G4>mRFP* (yellow arrows). NR: 65 -96hALH; Dissection: 96hALH. Scale bar = 20μm in (D’-E’’). (F) Quantification of the normalised (to Fed) Path-GFP intensity at the BBB in (D and E) (n = 8, 8). (G-H’’) Single-section images of Path-GFP expression in NBs (Mira, blue arrows) and CG (*repo-G4>RFP*, magenta arrows, distinguished from other glia based on location) in the *elav^C155^QF>QUAS-prosRi* tumour brains under Fed and NR. Scale bar = 20μm (I) Quantification of the normalised (to Fed) Path-GFP intensity of NBs and CG in (G-H’’) (n = 63, 45, 60, 35). (J-K) Maximum projection images of Path-GFP expression in *ey-FLP, act- G4> Ras^V12^, scribRi* eye disc tumours, marked by *UAS-RFP* under Fed and NR_._ (L) Quantification of Path-GFP intensity (CTCF, described in Methods) of circled tumour normalised to the tumour area in (M-N) (n = 6, 6). (J-L) are from the same experiment as Fig 1W-Z. Data information: ALH = hours after larvae hatching. NR: 72-120hALH; Dissection: 120hALH unless otherwise stated. Brain lobes are circled with yellow dashed lines. Scale bar = 50μm. Error bar represents SEM. In (C): unpaired t-test, (ns) P = 0.6740. In (F): unpaired t-test, (*) P = 0.0144. In (I): Mann–Whitney test, (*) P = 0.0240; Mann–Whitney test, (****) P < 0.0001. In (L): Mann–Whitney test, (ns) P = 0.1320.

**S7 Fig.**
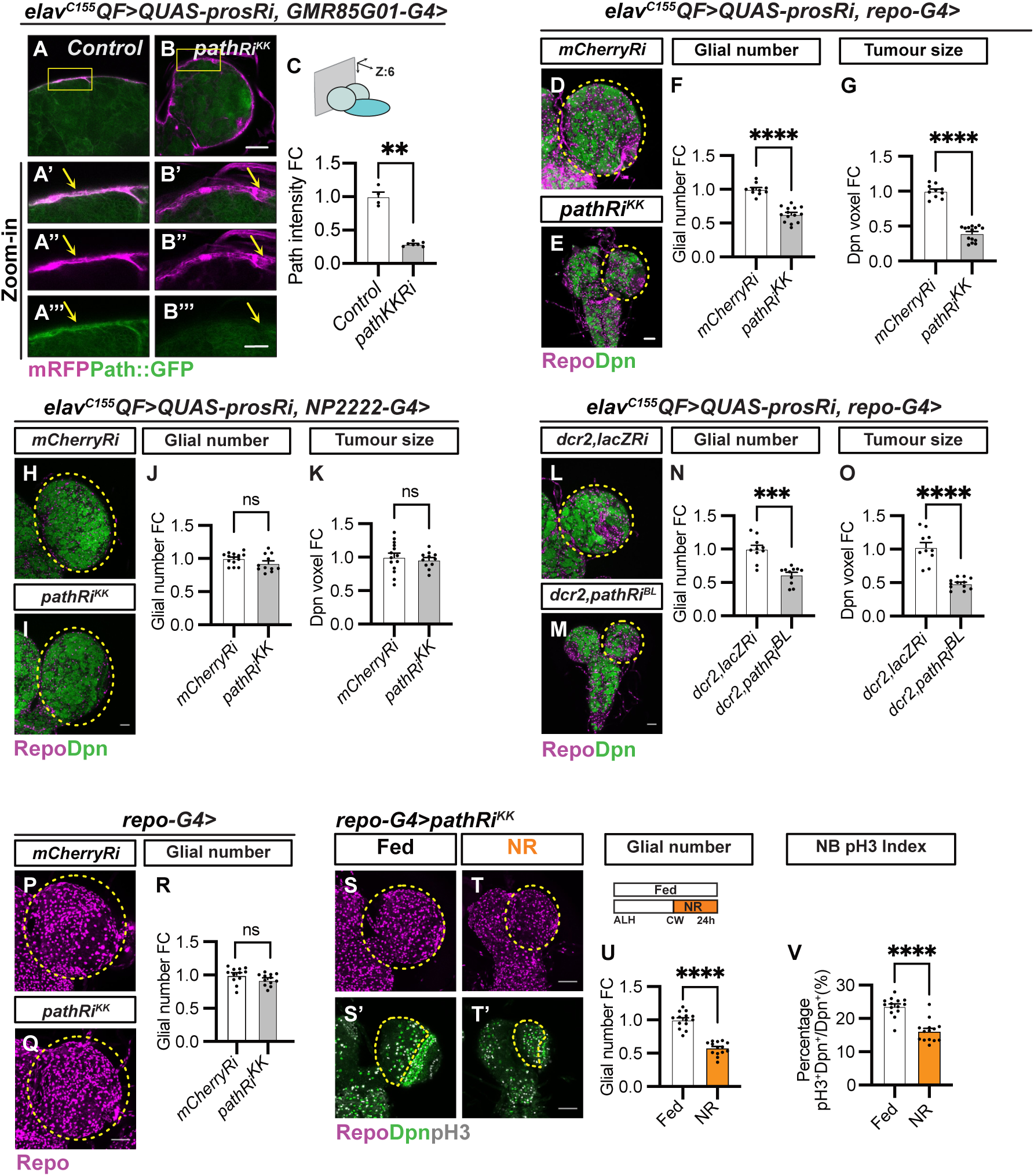
Characterisation of the role of Path in tumour glia and the wild-type glia (related to Fig 4). (A-B) Single-section (A and B) and zoomed-in images (A’-B’’’) of Path-GFP expression (yellow arrows) in *elav^C155^QF>QUAS-prosRi* tumour brains, where *pathRi^KK^* was overexpressed in PG using *GMR85G01-G4*, compared with *control* (CyOYFP siblings) at 96hALH. Scale bar = 20μm in (A’-B’’’) (C) Quantification of the normalised (to *control*) Path-GFP intensity at the PG membrane in (A-B) (n = 4, 7). (D-E) Single section images of *elav^C155^QF>QUAS-prosRi* tumour brains, where *mCherryRi* and *pathRi^KK^* were overexpressed in glia using *repo-G4*. Glial cells are stained with Repo, and NBs with Dpn. (F-G) Quantifications of the normalised (to *mCherryRi*) glial number (F) and Dpn voxels (G) of each circled brain lobe in (D-E) (n = 10, 14). (H-I) Single section images of *elav^C155^QF>QUAS-prosRi* tumour brains with *pathRi^KK^* (I) overexpressed specifically in CG using *NP2222-G4*, compared to *mCherryRi* (H) at 120hALH. Glia: Repo; NBs: Dpn. (J-K) Quantifications of the normalised (to *mCherryRi*) glial number (J) and Dpn voxels (K) of each circled brain lobe in (H-I) (n = 14, 11). (L-M) Single section images of *elav^C155^QF>QUAS-prosRi* tumour brains with an independent *pathRi* co-overexpressed with *dcr2* (M) in glia using *repo-G4*, compared to *dcr2, lacZRi* (L) at 120hALH. Glia: Repo; NBs: Dpn. (N-O) Quantifications of the normalised (to *dcr2, lacZRi*) glial number (N) and Dpn voxels (O) of each circled brain lobe in (L-M) (n = 10, 11). (P-Q) Maximum projection images of wild-type brain lobes with *pathRi^KK^* (Q) overexpressed in glia using *repo-G4*, compared to *mCherryRi* (P) at 96hALH. Glia: Repo. (R) Quantifications of the normalised (to *mCherryRi*) glial number of each circled brain lobe in (T-U) (n = 12, 12). (S-T’) Wild- type brain lobes where *pathRi^KK^*was overexpressed in glia using *repo-G4* under Fed and NR. Glial cells are stained with Repo in (S and T, maximum projection), Dpn and pH3 in (S’ and T’, single sections). CBs are circled by yellow dashed lines. NR: 65-89hALH; Dissection: 89hALH. (U) Quantification of the normalised (to Fed) glial number of each circled brain lobes in (S and T) (n = 14, 14). (V) Quantification of the percentage of NBs undergoing mitosis (pH3^+^Dpn^+^) in the CB of (S’ and T’) (n = 14, 14). Data information: ALH = hours after larvae hatching. Brain lobes are circled with yellow dashed lines. Scale bar = 50μm. Error bar represents SEM. In (C): One-way ANOVA, (**) P = 0.0015. In (F): One-way ANOVA, (****) P < 0.0001. In (G): Kruskal-Wallis test, (****) P < 0.0001. In (J): unpaired t-test, (ns) P = 0.1049. In (K): unpaired t-test, (ns) P = 0.6000. In (N): Mann–Whitney test, (***) P = 0.0001. In (O): Welch’s t-test, (****) P < 0.0001. In (R): unpaired t-test, (ns) P = 0.1245. In (U): unpaired t-test, (****) P < 0.0001. In (V): unpaired t- test, (****) P < 0.0001.

**S8 Fig.**
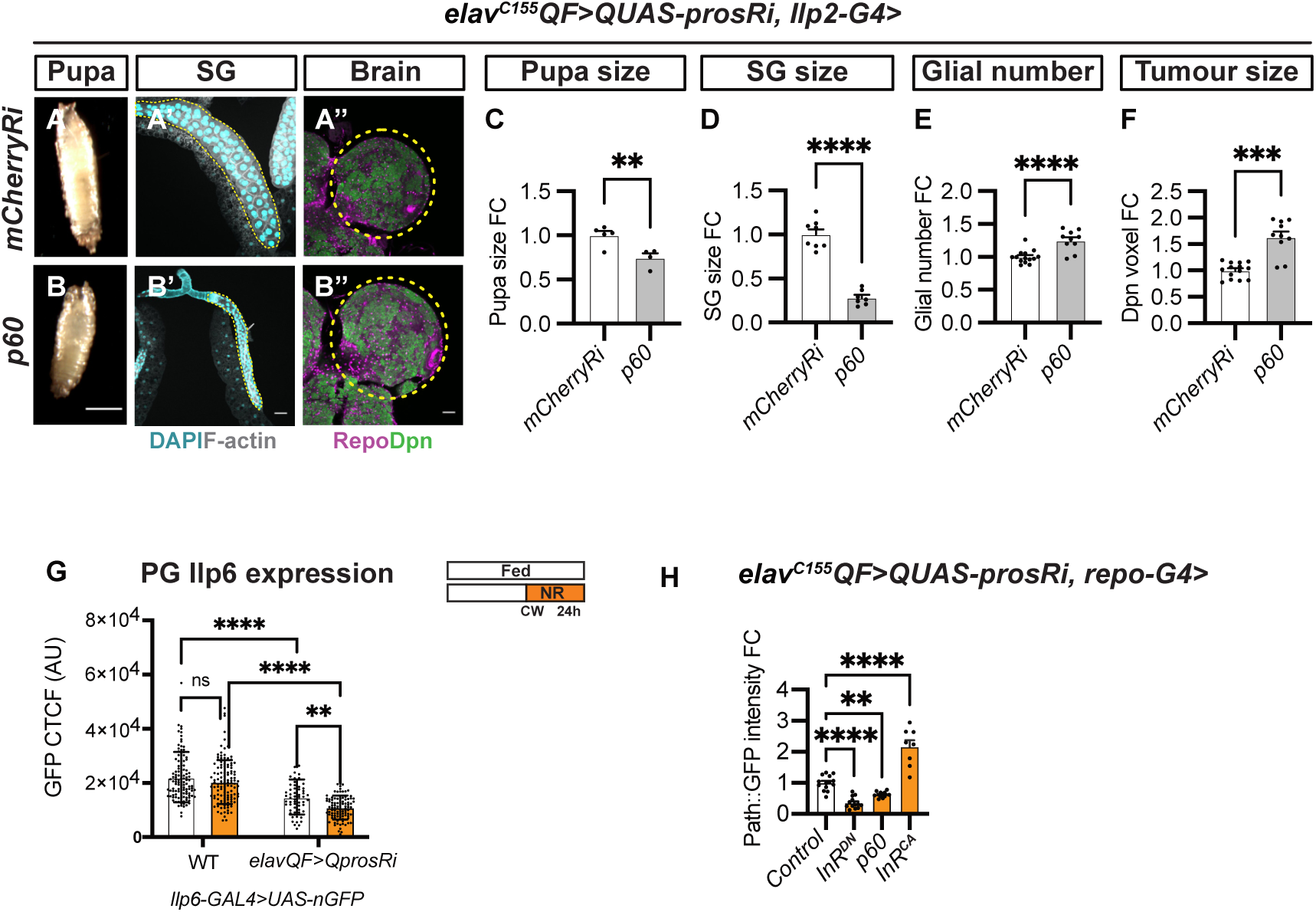
The glial PI3K pathway is not regulated by systemic Ilps (related to Fig. 6). (A-B) Pupa, salivary glands (SG, stained with Phalloidin and DAPI) and brains (stained with Repo and Dpn) in *elav^C155^QF>QUAS-prosRi* tumour-bearing animals, where *p60* was overexpressed in insulin-producing cells (IPCs) using *Ilp2-G4*, compared with *mCherryRi* at 120hALH. (A’) and (B’): maximum projection images; (A’’) and (B’’): single section images; Scale bar = 1mm in (A and B). (C) Quantification of the normalised (to *mCherryRi*) pupal size (area) in (A and B) (n = 5, 4). (D) Quantification of the normalised (to *mCherryRi*) SG size (area) in (A’ and B’) (n = 8, 7). (E-F) Quantifications of the normalised (to *mCherryRi*) glial number (E) and Dpn voxels (F) of each circled brain lobe in (A’’ and B’’) (n = 14, 10). (G) Quantifications of the GFP intensity (CTCF) in PG nucleus in wild-type and *elav^C155^QF>QUAS-prosRi* tumour brains under fed and NR conditions. Wild-type and tumour-bearing animals were starved after CW (65hALH and 68hALH, respectively) (n = 120, 120, 60, 100 cells from 12, 12, 6, 10 brain lobes). (H): Quantification of Path-GFP expression at the surface glia of the *elav^C155^QF>QUAS-prosRi* tumour brains with *InR^DN^*, *p60* or *InR^CA^* overexpressed in glia using *repo-G4* at 120hALH (n = 14, 13, 12, 8). Data information: ALH = after larvae hatching. Brain lobes are circled with yellow dashed lines. Scale bar = 50μm. Error bar represents SEM. In (C): unpaired t-test, (**) P = 0.0098. In (D): unpaired t-test, (****) P < 0.0001. In (E): unpaired t-test, (****) P < 0.0001. In (F): Welch’s t-test, (***) P = 0.0001. In (G): Two-way ANOVA was used to analyse whether the effect of NR on PG Ilp6 expression is different in the tumour brains compared with wild-type brains (significance indicated by interaction P value). Interaction P = 0.1876. Multiple comparisons: WT fed vs NR: (ns) P = 0.2488; Tumour fed vs NR: (**) P = 0.0095; WT fed vs Tumour fed: (****) P < 0.0001; WT NR vs Tumour NR: (****) P < 0.0001. In (H): One-way ANOVA, (****) P < 0.0001, (**) P = 0.0073, (****) P < 0.0001. In (M): One-way ANOVA, (****) P < 0.0001.

**S1 Table.**
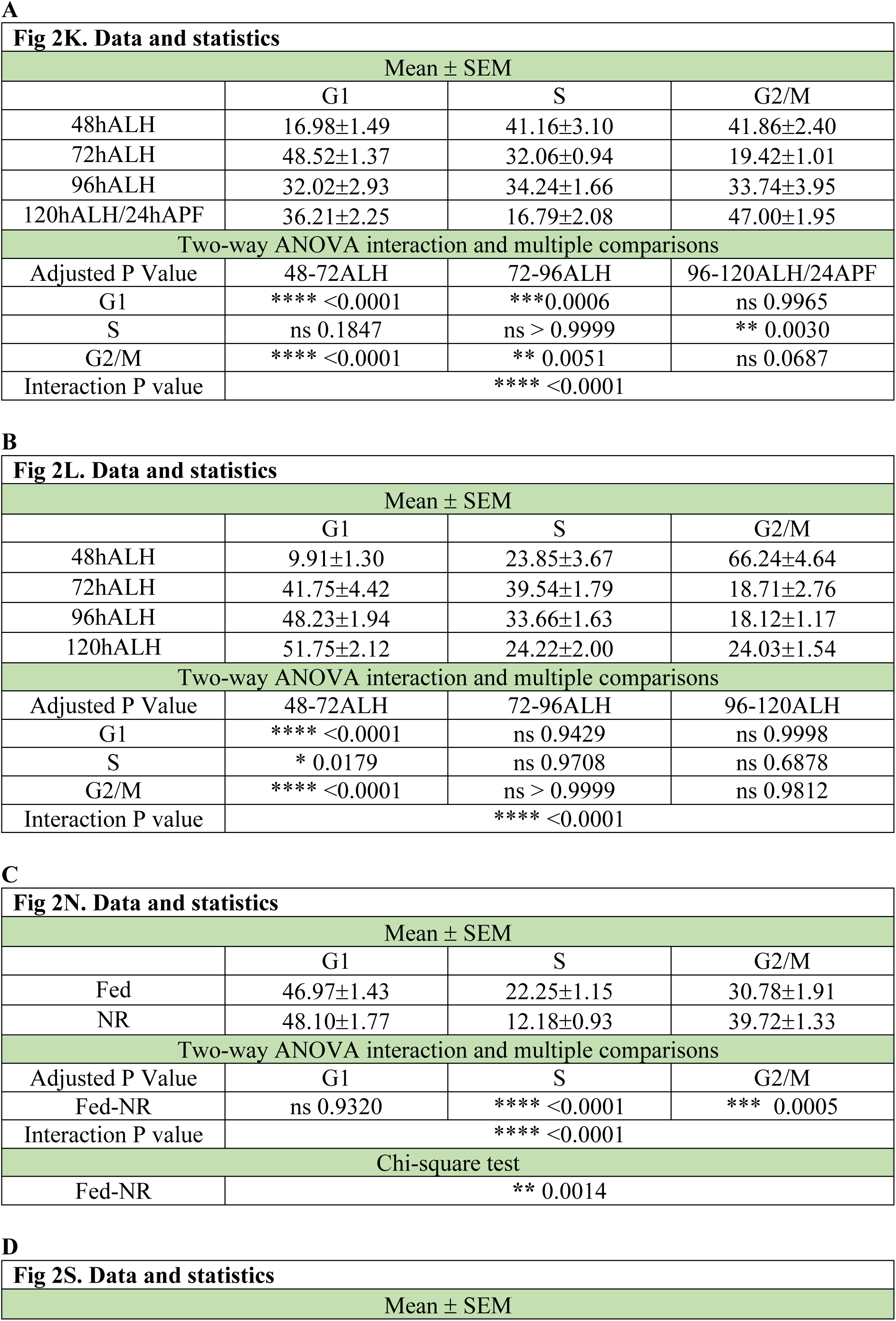
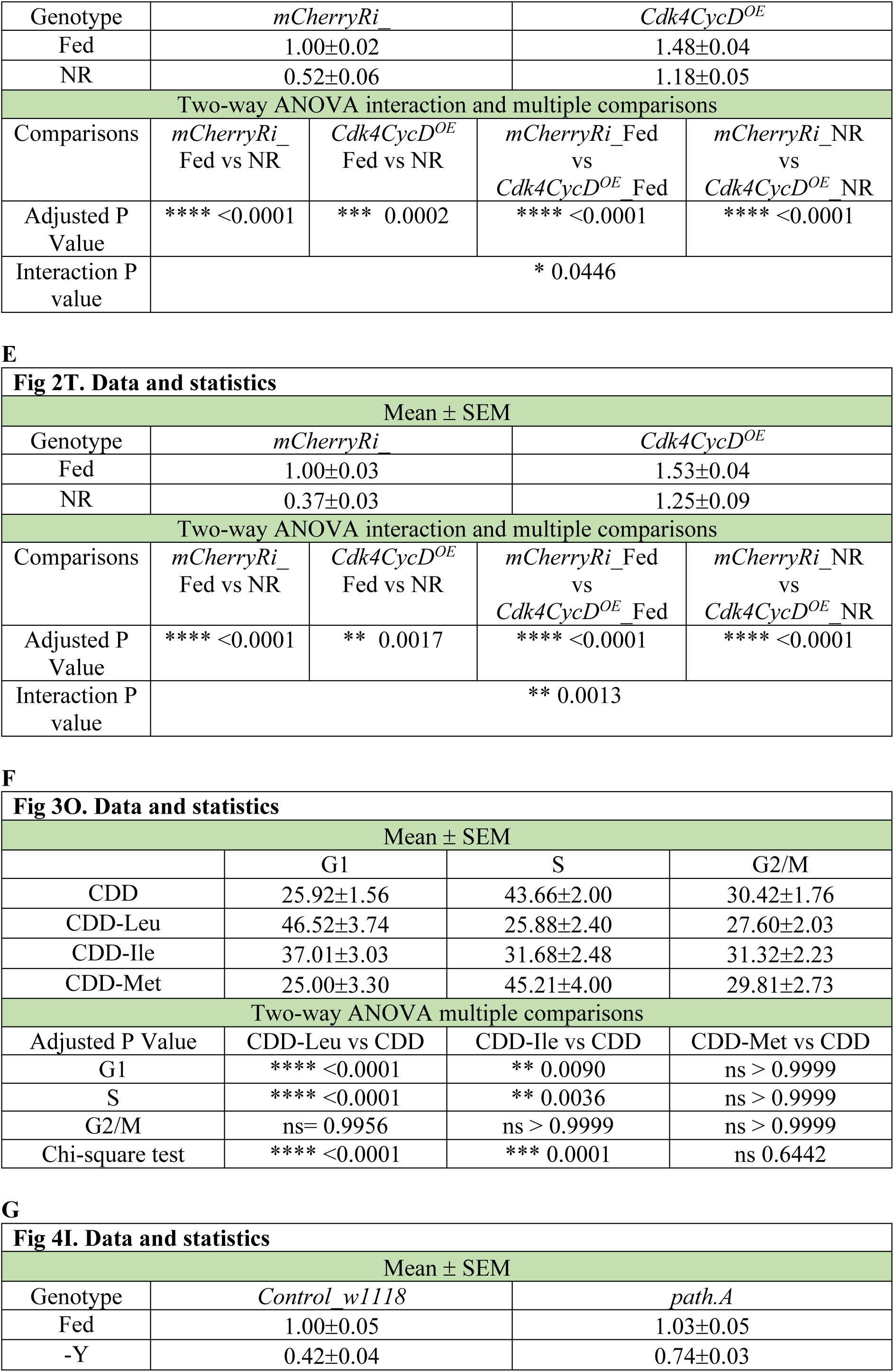
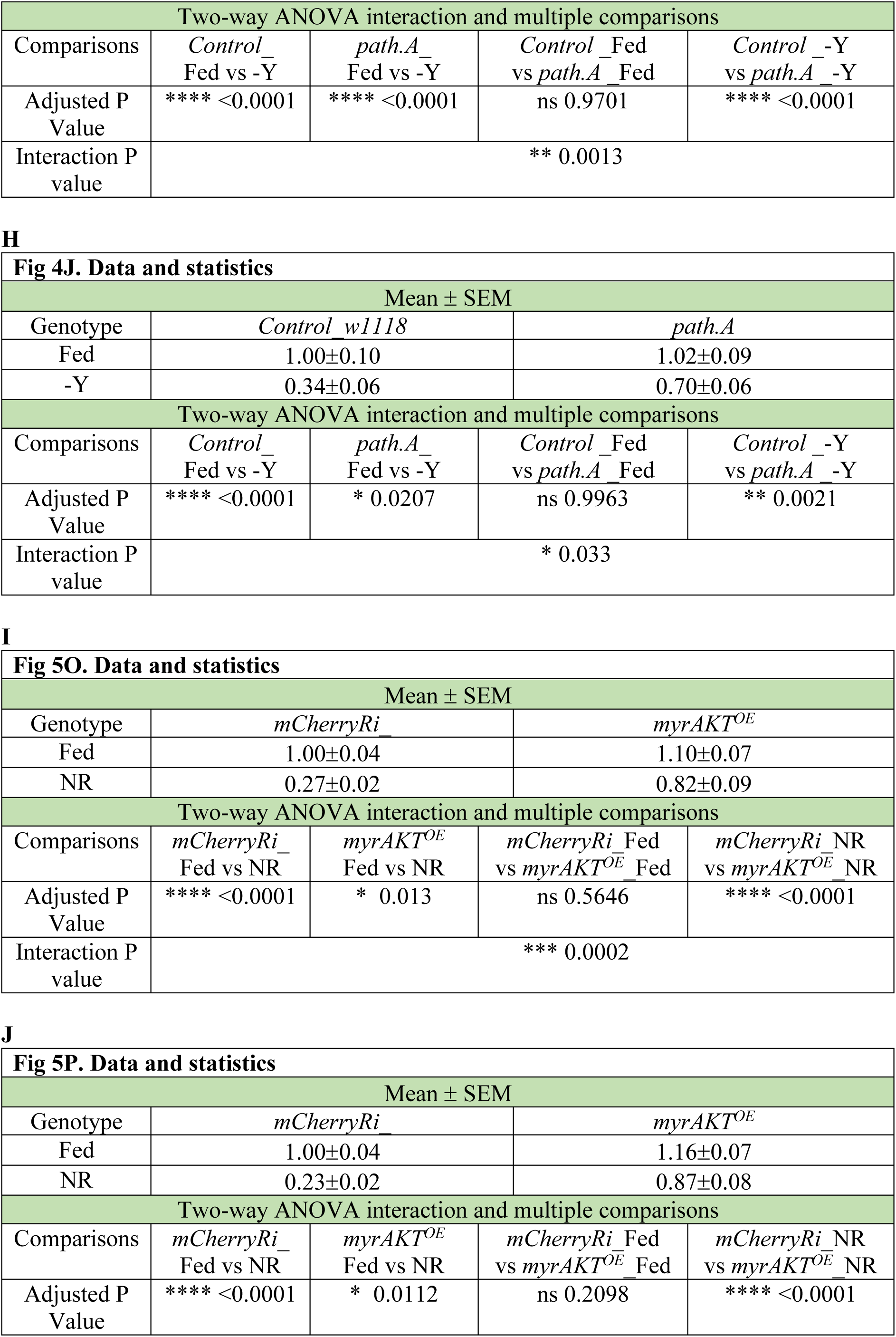
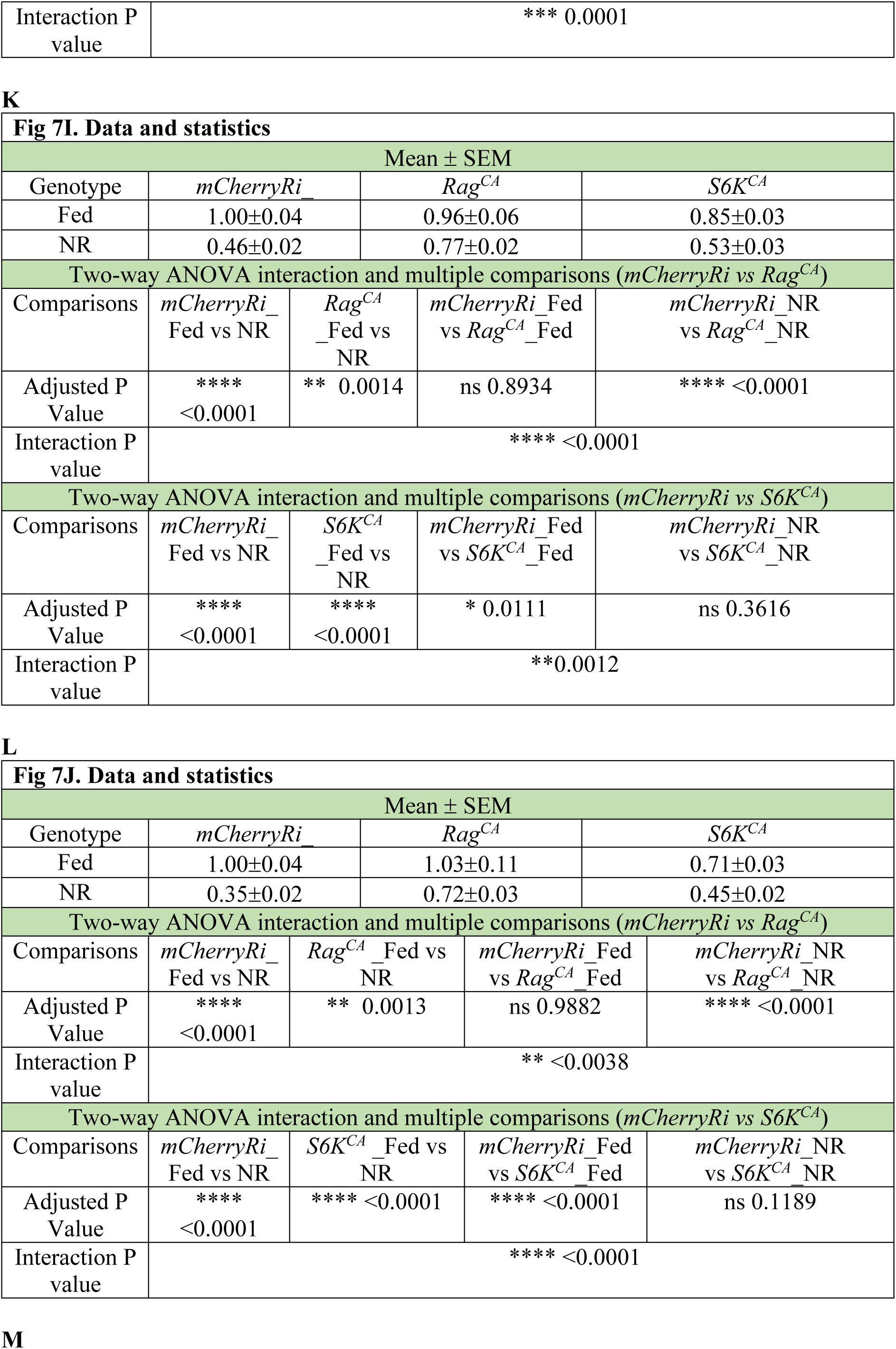
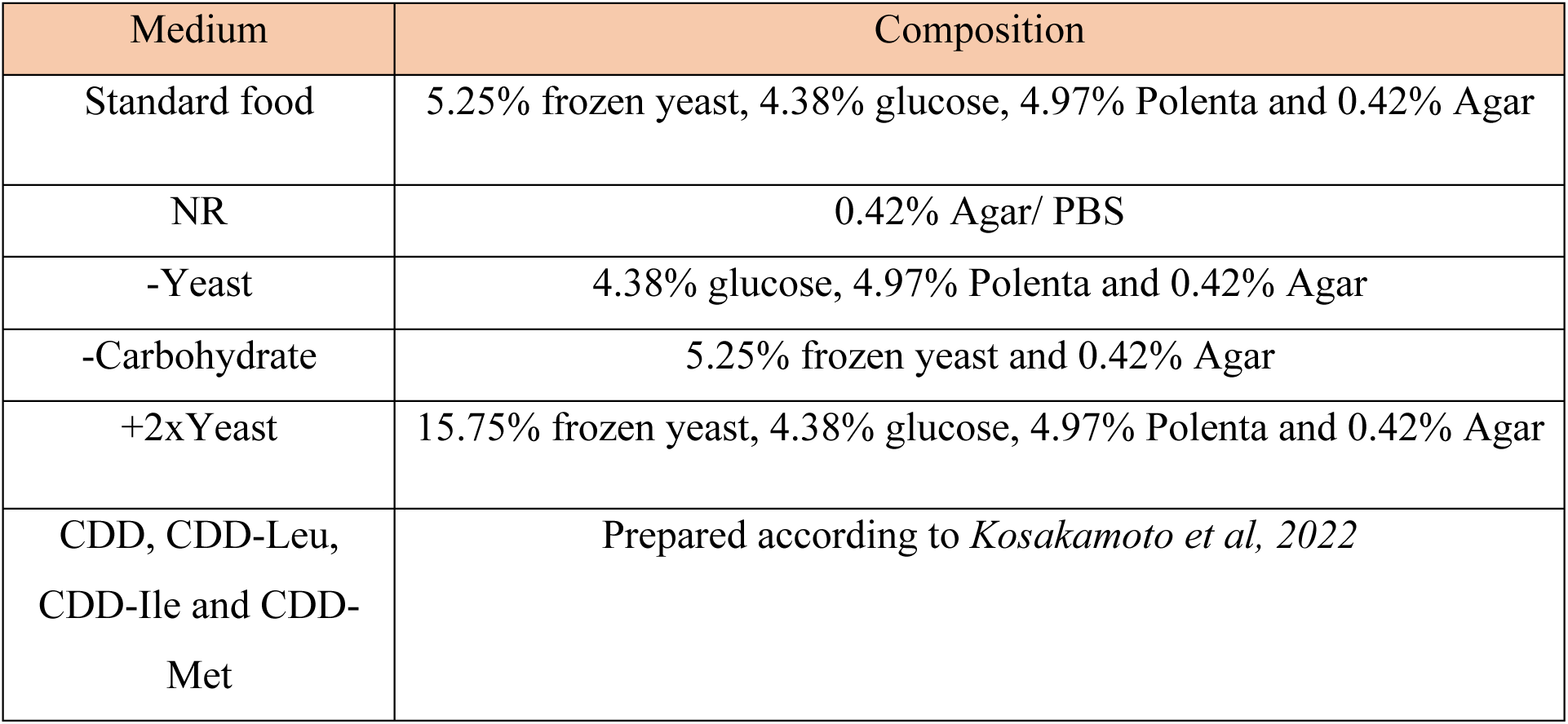
: This sheet includes Statistical data and the food recipe mentioned in methods. A B

| **Fig 2K. Data and statistics** |  |  |  |
| --- | --- | --- | --- |
| Mean ± SEM | | | |
|  | G1 | S | G2/M |
| 48hALH | 16.98±1.49 | 41.16±3.10 | 41.86±2.40 |
| 72hALH | 48.52±1.37 | 32.06±0.94 | 19.42±1.01 |
| 96hALH | 32.02±2.93 | 34.24±1.66 | 33.74±3.95 |
| 120hALH/24hAPF | 36.21±2.25 | 16.79±2.08 | 47.00±1.95 |
| Two-way ANOVA interaction and multiple comparisons |  |  |  |
| Adjusted P Value | 48-72ALH | 72-96ALH | 96-120ALH/24APF |
| G1 | **** <0.0001 | ***0.0006 | ns 0.9965 |
| S | ns 0.1847 | ns > 0.9999 | ** 0.0030 |
| G2/M | **** <0.0001 | ** 0.0051 | ns 0.0687 |
| Interaction P value | **** <0.0001 |  |  |

**B**
| **Fig 2L. Data and statistics** |  |  |  |
| --- | --- | --- | --- |
| Mean ± SEM | | | |
|  | G1 | S | G2/M |
| 48hALH | 9.91±1.30 | 23.85±3.67 | 66.24±4.64 |
| 72hALH | 41.75±4.42 | 39.54±1.79 | 18.71±2.76 |
| 96hALH | 48.23±1.94 | 33.66±1.63 | 18.12±1.17 |
| 120hALH | 51.75±2.12 | 24.22±2.00 | 24.03±1.54 |
| Two-way ANOVA interaction and multiple comparisons |  |  |  |
| Adjusted P Value | 48-72ALH | 72-96ALH | 96-120ALH |
| G1 | **** <0.0001 | ns 0.9429 | ns 0.9998 |
| S | * 0.0179 | ns 0.9708 | ns 0.6878 |
| G2/M | **** <0.0001 | ns > 0.9999 | ns 0.9812 |
| Interaction P value | **** <0.0001 |  |  |

**C**
| Fig 2N. Data and statistics |  |  |  |
| --- | --- | --- | --- |
| Mean ± SEM |  |  |  |
|  | G1 | S | G2/M |
| Fed | 46.97±1.43 | 22.25±1.15 | 30.78±1.91 |
| NR | 48.10±1.77 | 12.18±0.93 | 39.72±1.33 |
| Two-way ANOVA interaction and multiple comparisons |  |  |  |
| Adjusted P Value | G1 | S | G2/M |
| Fed-NR | ns 0.9320 | **** <0.0001 | *** 0.0005 |
| Interaction P value | **** <0.0001 |  |  |
| Chi-square test |  |  |  |
| Fed-NR | ** 0.0014 |  |  |

**D**
| **Fig 2S. Data and statistics** |  |  |  |
| --- | --- | --- | --- |
| Mean ± SEM | | | |

| Genotype | *mCherryRi* |  | *Cdk4CycD<sup>OE</sup>* |  |
| --- | --- | --- | --- | --- |
| Fed | 1.00±0.02 |  | 1.48±0.04 |  |
| NR | 0.52±0.06 |  | 1.18±0.05 |  |
| Two-way ANOVA interaction and multiple comparisons |  |  |  |  |
| Comparisons | *mCherryRi*_Fed vs NR | *Cdk4CycD<sup>OE</sup>*_Fed vs NR | *mCherryRi*_Fed vs *Cdk4CycD<sup>OE</sup>*_Fed | *mCherryRi*_NR vs *Cdk4CycD<sup>OE</sup>*_NR |
| Adjusted P Value | **** <0.0001 | *** 0.0002 | **** <0.0001 | **** <0.0001 |
| Interaction P value | * 0.0446 |  |  |  |

**E**
| Fig 2T. Data and statistics |  |  |  |  |
| --- | --- | --- | --- | --- |
| Mean ± SEM |  |  |  |  |
| Genotype | *mCherryRi* |  | *Cdk4CycD<sup>OE</sup>* |  |
| Fed | 1.00±0.03 |  | 1.53±0.04 |  |
| NR | 0.37±0.03 |  | 1.25±0.09 |  |
| Two-way ANOVA interaction and multiple comparisons |  |  |  |  |
| Comparisons | *mCherryRi*_Fed vs NR | *Cdk4CycD<sup>OE</sup>*_Fed vs NR | *mCherryRi*_Fed vs *Cdk4CycD<sup>OE</sup>*_Fed | *mCherryRi*_NR vs *Cdk4CycD<sup>OE</sup>*_NR |
| Adjusted P Value | **** <0.0001 | ** 0.0017 | **** <0.0001 | **** <0.0001 |
| Interaction P value | ** 0.0013 |  |  |  |

**F**
| Fig 3O. Data and statistics |  |  |  |
| --- | --- | --- | --- |
| Mean ± SEM |  |  |  |
|  | G1 | S | G2/M |
| CDD | 25.92±1.56 | 43.66±2.00 | 30.42±1.76 |
| CDD-Leu | 46.52±3.74 | 25.88±2.40 | 27.60±2.03 |
| CDD-Ile | 37.01±3.03 | 31.68±2.48 | 31.32±2.23 |
| CDD-Met | 25.00±3.30 | 45.21±4.00 | 29.81±2.73 |
| Two-way ANOVA multiple comparisons |  |  |  |
| Adjusted P Value | CDD-Leu vs CDD | CDD-Ile vs CDD | CDD-Met vs CDD |
| G1 | **** <0.0001 | ** 0.0090 | ns > 0.9999 |
| S | **** <0.0001 | ** 0.0036 | ns > 0.9999 |
| G2/M | ns= 0.9956 | ns > 0.9999 | ns > 0.9999 |
| Chi-square test | **** <0.0001 | *** 0.0001 | ns 0.6442 |

**G**
| Fig 4I. Data and statistics |  |  |
| --- | --- | --- |
| Mean ± SEM |  |  |
| Genotype | *Control w1118* | *path.A* |
| Fed | 1.00±0.05 | 1.03±0.05 |
| -Y | 0.42±0.04 | 0.74±0.03 |

| Two-way ANOVA interaction and multiple comparisons |  |  |  |  |
| --- | --- | --- | --- | --- |
| Comparisons | *Control*<sub>-</sub><br>Fed vs -Y | *path.A*<sub>-</sub><br>Fed vs -Y | *Control*<sub>-</sub> Fed<br>vs *path.A*<sub>-</sub> Fed | *Control*<sub>-</sub> -Y<br>vs *path.A*<sub>-</sub> -Y |
| Adjusted P Value | **** <0.0001 | **** <0.0001 | ns 0.9701 | **** <0.0001 |
| Interaction P value | ** 0.0013 |  |  |  |

**H**
| Fig 4J. Data and statistics |  |  |  |  |
| --- | --- | --- | --- | --- |
| Mean ± SEM |  |  |  |  |
| Genotype | *Control w1118* |  | *path.A* |  |
| Fed | 1.00±0.10 |  | 1.02±0.09 |  |
| -Y | 0.34±0.06 |  | 0.70±0.06 |  |
| Two-way ANOVA interaction and multiple comparisons |  |  |  |  |
| Comparisons | *Control*<br>Fed vs -Y | *path.A*<br>Fed vs -Y | *Control* Fed<br>vs *path.A* Fed | *Control* -Y<br>vs *path.A* -Y |
| Adjusted P Value | **** <0.0001 | * 0.0207 | ns 0.9963 | ** 0.0021 |
| Interaction P value | * 0.033 |  |  |  |

**I**
| Fig 5O. Data and statistics |  |  |  |  |
| --- | --- | --- | --- | --- |
| Mean ± SEM |  |  |  |  |
| Genotype | *mCherryRi*<sub>-</sub> |  | *myrAKT<sup>OE</sup>* |  |
| Fed | 1.00±0.04 |  | 1.10±0.07 |  |
| NR | 0.27±0.02 |  | 0.82±0.09 |  |
| Two-way ANOVA interaction and multiple comparisons |  |  |  |  |
| Comparisons | *mCherryRi*<sub>-</sub><br>Fed vs NR | *myrAKT<sup>OE</sup>*<br>Fed vs NR | *mCherryRi* Fed<br>vs *myrAKT<sup>OE</sup>* Fed | *mCherryRi* NR<br>vs *myrAKT<sup>OE</sup>* NR |
| Adjusted P Value | **** <0.0001 | * 0.013 | ns 0.5646 | **** <0.0001 |
| Interaction P value | *** 0.0002 |  |  |  |

**J**
| Fig 5P. Data and statistics |  |  |  |  |
| --- | --- | --- | --- | --- |
| Mean ± SEM |  |  |  |  |
| Genotype | *mCherryRi* |  | *myrAKT<sup>OE</sup>* |  |
| Fed | 1.00±0.04 |  | 1.16±0.07 |  |
| NR | 0.23±0.02 |  | 0.87±0.08 |  |
| Two-way ANOVA interaction and multiple comparisons |  |  |  |  |
| Comparisons | *mCherryRi*<br>Fed vs NR | *myrAKT<sup>OE</sup>*<br>Fed vs NR | *mCherryRi* Fed<br>vs *myrAKT<sup>OE</sup>* Fed | *mCherryRi* NR<br>vs *myrAKT<sup>OE</sup>* NR |
| Adjusted P Value | **** <0.0001 | * 0.0112 | ns 0.2098 | **** <0.0001 |

|  |  |
| --- | --- |
| Interaction P value | *** 0.0001 |

**K**
| Fig 7I. Data and statistics |  |  |  |  |
| --- | --- | --- | --- | --- |
| Mean ± SEM |  |  |  |  |
| Genotype | *mCherryRi*_ | *Rag<sup>CA</sup>* | *S6K<sup>CA</sup>* |  |
| Fed | 1.00±0.04 | 0.96±0.06 | 0.85±0.03 |  |
| NR | 0.46±0.02 | 0.77±0.02 | 0.53±0.03 |  |
| Two-way ANOVA interaction and multiple comparisons (*mCherryRi* vs *Rag<sup>CA</sup>*) |  |  |  |  |
| Comparisons | *mCherryRi*_Fed vs NR | *Rag<sup>CA</sup>*_Fed vs NR | *mCherryRi*_Fed vs *Rag<sup>CA</sup>*_Fed | *mCherryRi*_NR vs *Rag<sup>CA</sup>*_NR |
| Adjusted P Value | **** <0.0001 | ** 0.0014 | ns 0.8934 | **** <0.0001 |
| Interaction P value | **** <0.0001 |  |  |  |
| Two-way ANOVA interaction and multiple comparisons (*mCherryRi* vs *S6K<sup>CA</sup>*) |  |  |  |  |
| Comparisons | *mCherryRi*_Fed vs NR | *S6K<sup>CA</sup>*_Fed vs NR | *mCherryRi*_Fed vs *S6K<sup>CA</sup>*_Fed | *mCherryRi*_NR vs *S6K<sup>CA</sup>*_NR |
| Adjusted P Value | **** <0.0001 | **** <0.0001 | * 0.0111 | ns 0.3616 |
| Interaction P value | **0.0012 |  |  |  |

**L**
| Fig 7J. Data and statistics |  |  |  |  |
| --- | --- | --- | --- | --- |
| Mean ± SEM |  |  |  |  |
| Genotype | *mCherryRi* | *Rag<sup>CA</sup>* | *S6K<sup>CA</sup>* |  |
| Fed | 1.00±0.04 | 1.03±0.11 | 0.71±0.03 |  |
| NR | 0.35±0.02 | 0.72±0.03 | 0.45±0.02 |  |
| Two-way ANOVA interaction and multiple comparisons (*mCherryRi* vs *Rag<sup>CA</sup>*) |  |  |  |  |
| Comparisons | *mCherryRi*<br>Fed vs NR | *Rag<sup>CA</sup>* Fed vs<br>NR | *mCherryRi* Fed<br>vs *Rag<sup>CA</sup>* Fed | *mCherryRi* NR<br>vs *Rag<sup>CA</sup>* NR |
| Adjusted P<br>Value | ****<br><0.0001 | ** 0.0013 | ns 0.9882 | **** <0.0001 |
| Interaction P<br>value | ** <0.0038 |  |  |  |
| Two-way ANOVA interaction and multiple comparisons (*mCherryRi* vs *S6K<sup>CA</sup>*) |  |  |  |  |
| Comparisons | *mCherryRi*<br>Fed vs NR | *S6K<sup>CA</sup>* Fed vs<br>NR | *mCherryRi* Fed<br>vs *S6K<sup>CA</sup>* Fed | *mCherryRi* NR<br>vs *S6K<sup>CA</sup>* NR |
| Adjusted P<br>Value | ****<br><0.0001 | **** <0.0001 | **** <0.0001 | ns 0.1189 |
| Interaction P<br>value | **** <0.0001 |  |  |  |

**M**
| Medium | Composition |
| --- | --- |
| Standard food | 5.25% frozen yeast, 4.38% glucose, 4.97% Polenta and 0.42% Agar |
| NR | 0.42% Agar/ PBS |
| -Yeast | 4.38% glucose, 4.97% Polenta and 0.42% Agar |
| -Carbohydrate | 5.25% frozen yeast and 0.42% Agar |
| +2xYeast | 15.75% frozen yeast, 4.38% glucose, 4.97% Polenta and 0.42% Agar |
| CDD, CDD-Leu,<br>CDD-Ile and CDD-Met | Prepared according to *Kosakamoto et al, 2022* |

S1 Data: Excel spreadsheet showing genes significantly upregulated and downregulated in tumour brains under NR compared to Fed (False Discovery Rate (FDR) ≤ 0.05, Fold Change (FC) ≥ 1.5).

S2 Data: Excel spreadsheet showing the expression of annotated transporters (using GLAD) in tumour brains under fed and NR conditions (False discovery rate, FDR<0.05; Fold change (FC) >1.5)

## Reference

1. Faubert B, Solmonson A, DeBerardinis RJ. Metabolic reprogramming and cancer progression. Science. 2020;368(6487).

2. Froldi F, Pachnis P, Szuperak M, Costas O, Fernando T, Gould AP, et al. Histidine is selectively required for the growth of Myc-dependent dedifferentiation tumours in the Drosophila CNS. EMBO J. 2019;38(7).

3. Kalaany NY, Sabatini DM. Author Correction: Tumours with PI3K activation are resistant to dietary restriction. Nature. 2020;581(7807):E2.

4. Mukherjee P, El-Abbadi MM, Kasperzyk JL, Ranes MK, Seyfried TN. Dietary restriction reduces angiogenesis and growth in an orthotopic mouse brain tumour model. Br J Cancer. 2002;86(10):1615–21.

5. Upadhyayula PS, Higgins DM, Mela A, Banu M, Dovas A, Zandkarimi F, et al. Dietary restriction of cysteine and methionine sensitizes gliomas to ferroptosis and induces alterations in energetic metabolism. Nat Commun. 2023;14(1):1187.

6. Bielecka-Wajdman AM, Ludyga T, Smyk D, Smyk W, Mularska M, Swiderek P, et al. Glucose Influences the Response of Glioblastoma Cells to Temozolomide and Dexamethasone. Cancer Control. 2022;29:10732748221075468.

7. Bao Z, Chen K, Krepel S, Tang P, Gong W, Zhang M, et al. High Glucose Promotes Human Glioblastoma Cell Growth by Increasing the Expression and Function of Chemoattractant and Growth Factor Receptors. Transl Oncol. 2019;12(9):1155–63.

8. Nguyen PK, Cheng LY. Non-autonomous regulation of neurogenesis by extrinsic cues: a Drosophila perspective. Oxf Open Neurosci. 2022;1:kvac004.

9. Ding WY, Huang J, Wang H. Waking up quiescent neural stem cells: Molecular mechanisms and implications in neurodevelopmental disorders. PLoS Genet. 2020;16(4):e1008653.

10. Rujano MA, Briand D, Ethelic B, Marc J, Speder P. An interplay between cellular growth and atypical fusion defines morphogenesis of a modular glial niche in Drosophila. Nat Commun. 2022;13(1):4999.

11. Dumstrei K, Wang F, Hartenstein V. Role of DE-cadherin in neuroblast proliferation, neural morphogenesis, and axon tract formation in Drosophila larval brain development. J Neurosci. 2003;23(8):3325–35.

12. Pereanu W, Shy D, Hartenstein V. Morphogenesis and proliferation of the larval brain glia in Drosophila. Dev Biol. 2005;283(1):191–203.

13. Speder P, Brand AH. Systemic and local cues drive neural stem cell niche remodelling during neurogenesis in Drosophila. Elife. 2018;7.

14. Yuan X, Sipe CW, Suzawa M, Bland ML, Siegrist SE. Dilp-2-mediated PI3-kinase activation coordinates reactivation of quiescent neuroblasts with growth of their glial stem cell niche. PLoS Biol. 2020;18(5):e3000721.

15. Stork T, Engelen D, Krudewig A, Silies M, Bainton RJ, Klambt C. Organization and function of the blood-brain barrier in Drosophila. J Neurosci. 2008;28(3):587–97.

16. Limmer S, Weiler A, Volkenhoff A, Babatz F, Klambt C. The Drosophila blood-brain barrier: development and function of a glial endothelium. Front Neurosci. 2014;8:365.

17. Contreras EG, Klambt C. The Drosophila blood-brain barrier emerges as a model for understanding human brain diseases. Neurobiol Dis. 2023;180:106071.

18. Mayer F, Mayer N, Chinn L, Pinsonneault RL, Kroetz D, Bainton RJ. Evolutionary conservation of vertebrate blood-brain barrier chemoprotective mechanisms in Drosophila. J Neurosci. 2009;29(11):3538–50.

19. McMullen E, Weiler A, Becker HM, Schirmeier S. Plasticity of Carbohydrate Transport at the Blood-Brain Barrier. Front Behav Neurosci. 2020;14:612430.

20. Hindle SJ, Munji RN, Dolghih E, Gaskins G, Orng S, Ishimoto H, et al. Evolutionarily Conserved Roles for Blood-Brain Barrier Xenobiotic Transporters in Endogenous Steroid Partitioning and Behavior. Cell Rep. 2017;21(5):1304–16.

21. Li F, Artiushin G, Sehgal A. Modulation of sleep by trafficking of lipids through the Drosophila blood-brain barrier. Elife. 2023;12.

22. Choksi SP, Southall TD, Bossing T, Edoff K, de Wit E, Fischer BE, et al. Prospero acts as a binary switch between self-renewal and differentiation in Drosophila neural stem cells. Dev Cell. 2006;11(6):775–89.

23. Caussinus E, Gonzalez C. Induction of tumor growth by altered stem-cell asymmetric division in Drosophila melanogaster. Nat Genet. 2005;37(10):1125–9.

24. Cheng LY, Bailey AP, Leevers SJ, Ragan TJ, Driscoll PC, Gould AP. Anaplastic lymphoma kinase spares organ growth during nutrient restriction in Drosophila. Cell. 2011;146(3):435–47.

25. Feng S, Zacharioudaki E, Millen K, Bray SJ. The SLC36 transporter Pathetic is required for neural stem cell proliferation and for brain growth under nutrition restriction. Neural Dev. 2020;15(1):10.

26. Hertenstein H, McMullen E, Weiler A, Volkenhoff A, Becker HM, Schirmeier S. Starvation-induced regulation of carbohydrate transport at the blood-brain barrier is TGF- beta-signaling dependent. Elife. 2021;10.

27. Alvarez-Ochoa E, Dong Q, Truong H, Cheng LY. Tumour-derived gliogenesis sustains dedifferentiation-dependent tumour growth in the *Drosophila* CNS. bioRxiv. 2024:2024.12.23.630170.

28. Clark JM, Gibbs AG. Starvation selection reduces and delays larval ecdysone production and signaling. J Exp Biol. 2023;226(18).

29. Lee CY, Robinson KJ, Doe CQ. Lgl, Pins and aPKC regulate neuroblast self-renewal versus differentiation. Nature. 2006;439(7076):594-8.

30. Bowman SK, Rolland V, Betschinger J, Kinsey KA, Emery G, Knoblich JA. The tumor suppressors Brat and Numb regulate transit-amplifying neuroblast lineages in Drosophila. Dev Cell. 2008;14(4):535–46.

31. Avet-Rochex A, Kaul AK, Gatt AP, McNeill H, Bateman JM. Concerted control of gliogenesis by InR/TOR and FGF signalling in the Drosophila post-embryonic brain. Development. 2012;139(15):2763–72.

32. Brumby AM, Richardson HE. scribble mutants cooperate with oncogenic Ras or Notch to cause neoplastic overgrowth in Drosophila. EMBO J. 2003;22(21):5769–79.

33. Contreras EG, Glavic A, Brand AH, Sierralta JA. The Serine Protease Homolog, Scarface, Is Sensitive to Nutrient Availability and Modulates the Development of the Drosophila Blood-Brain Barrier. J Neurosci. 2021;41(30):6430–48.

34. Zielke N, Korzelius J, van Straaten M, Bender K, Schuhknecht GFP, Dutta D, et al. Fly-FUCCI: A versatile tool for studying cell proliferation in complex tissues. Cell Rep. 2014;7(2):588–98.

35. Kremer MC, Jung C, Batelli S, Rubin GM, Gaul U. The glia of the adult Drosophila nervous system. Glia. 2017;65(4):606–38.

36. Rattanapornsompong K, Khattiya J, Phannasil P, Phaonakrop N, Roytrakul S, Jitrapakdee S, et al. Impaired G2/M cell cycle arrest induces apoptosis in pyruvate carboxylase knockdown MDA-MB-231 cells. Biochem Biophys Rep. 2021;25:100903.

37. Maniere G, Ziegler AB, Geillon F, Featherstone DE, Grosjean Y. Direct Sensing of Nutrients via a LAT1-like Transporter in Drosophila Insulin-Producing Cells. Cell Rep. 2016;17(1):137–48.

38. Martin JF, Hersperger E, Simcox A, Shearn A. minidiscs encodes a putative amino acid transporter subunit required non-autonomously for imaginal cell proliferation. Mech Dev. 2000;92(2):155–67.

39. Reynolds B, Roversi P, Laynes R, Kazi S, Boyd CA, Goberdhan DC. Drosophila expresses a CD98 transporter with an evolutionarily conserved structure and amino acid- transport properties. Biochem J. 2009;420(3):363–72.

40. Kosakamoto H, Okamoto N, Aikawa H, Sugiura Y, Suematsu M, Niwa R, et al. Sensing of the non-essential amino acid tyrosine governs the response to protein restriction in Drosophila. Nat Metab. 2022;4(7):944–59.

41. Piper MD, Blanc E, Leitao-Goncalves R, Yang M, He X, Linford NJ, et al. A holidic medium for Drosophila melanogaster. Nat Methods. 2014;11(1):100–5.

42. Sowers ML, Sowers LC. Glioblastoma and Methionine Addiction. Int J Mol Sci. 2022;23(13).

43. Neinast M, Murashige D, Arany Z. Branched Chain Amino Acids. Annu Rev Physiol. 2019;81:139–64.

44. Marygold SJ. The alpha-ketoacid dehydrogenase complexes of Drosophila melanogaster. MicroPubl Biol. 2024;2024.

45. Lin WY, Williams CR, Yan C, Parrish JZ. Functions of the SLC36 transporter Pathetic in growth control. Fly (Austin). 2015;9(3):99–106.

46. Awasaki T, Lai SL, Ito K, Lee T. Organization and postembryonic development of glial cells in the adult central brain of Drosophila. J Neurosci. 2008;28(51):13742–53.

47. Lin WY, Williams C, Yan C, Koledachkina T, Luedke K, Dalton J, et al. The SLC36 transporter Pathetic is required for extreme dendrite growth in Drosophila sensory neurons. Genes Dev. 2015;29(11):1120–35.

48. Pillai SM, Meredith D. SLC36A4 (hPAT4) is a high affinity amino acid transporter when expressed in Xenopus laevis oocytes. J Biol Chem. 2011;286(4):2455–60.

49. Goberdhan DC, Meredith D, Boyd CA, Wilson C. PAT-related amino acid transporters regulate growth via a novel mechanism that does not require bulk transport of amino acids. Development. 2005;132(10):2365–75.

50. Britton JS, Lockwood WK, Li L, Cohen SM, Edgar BA. Drosophila’s insulin/PI3- kinase pathway coordinates cellular metabolism with nutritional conditions. Dev Cell. 2002;2(2):239–49.

51. Brunet A, Bonni A, Zigmond MJ, Lin MZ, Juo P, Hu LS, et al. Akt promotes cell survival by phosphorylating and inhibiting a Forkhead transcription factor. Cell. 1999;96(6):857–68.

52. Ikeya T, Galic M, Belawat P, Nairz K, Hafen E. Nutrient-dependent expression of insulin-like peptides from neuroendocrine cells in the CNS contributes to growth regulation in Drosophila. Curr Biol. 2002;12(15):1293–300.

53. Rulifson EJ, Kim SK, Nusse R. Ablation of insulin-producing neurons in flies: growth and diabetic phenotypes. Science. 2002;296(5570):1118-20.

54. Okamoto N, Yamanaka N, Yagi Y, Nishida Y, Kataoka H, O’Connor MB, et al. A fat body-derived IGF-like peptide regulates postfeeding growth in Drosophila. Dev Cell. 2009;17(6):885–91.

55. Slaidina M, Delanoue R, Gronke S, Partridge L, Leopold P. A Drosophila insulin-like peptide promotes growth during nonfeeding states. Dev Cell. 2009;17(6):874–84.

56. Sousa-Nunes R, Yee LL, Gould AP. Fat cells reactivate quiescent neuroblasts via TOR and glial insulin relays in Drosophila. Nature. 2011;471(7339):508-12.

57. Bai H, Kang P, Tatar M. Drosophila insulin-like peptide-6 (dilp6) expression from fat body extends lifespan and represses secretion of Drosophila insulin-like peptide-2 from the brain. Aging Cell. 2012;11(6):978–85.

58. Newton H, Wang YF, Camplese L, Mokochinski JB, Kramer HB, Brown AEX, et al. Systemic muscle wasting and coordinated tumour response drive tumourigenesis. Nat Commun. 2020;11(1):4653.

59. Shang P, Valapala M, Grebe R, Hose S, Ghosh S, Bhutto IA, et al. The amino acid transporter SLC36A4 regulates the amino acid pool in retinal pigmented epithelial cells and mediates the mechanistic target of rapamycin, complex 1 signaling. Aging Cell. 2017;16(2):349–59.

60. Schwitalla S, Fingerle AA, Cammareri P, Nebelsiek T, Goktuna SI, Ziegler PK, et al. Intestinal tumorigenesis initiated by dedifferentiation and acquisition of stem-cell-like properties. Cell. 2013;152(1-2):25–38.

61. Friedmann-Morvinski D, Bushong EA, Ke E, Soda Y, Marumoto T, Singer O, et al. Dedifferentiation of neurons and astrocytes by oncogenes can induce gliomas in mice. Science. 2012;338(6110):1080-4.

62. Stine ZE, Schug ZT, Salvino JM, Dang CV. Targeting cancer metabolism in the era of precision oncology. Nat Rev Drug Discov. 2022;21(2):141–62.

63. Lobel GP, Jiang Y, Simon MC. Tumor microenvironmental nutrients, cellular responses, and cancer. Cell Chem Biol. 2023;30(9):1015–32.

64. Thwaites DT, Anderson CM. The SLC36 family of proton-coupled amino acid transporters and their potential role in drug transport. Br J Pharmacol. 2011;164(7):1802–16.

65. Heublein S, Kazi S, Ogmundsdottir MH, Attwood EV, Kala S, Boyd CA, et al. Proton-assisted amino-acid transporters are conserved regulators of proliferation and amino- acid-dependent mTORC1 activation. Oncogene. 2010;29(28):4068–79.

66. Yilmaz OH, Katajisto P, Lamming DW, Gultekin Y, Bauer-Rowe KE, Sengupta S, et al. mTORC1 in the Paneth cell niche couples intestinal stem-cell function to calorie intake. Nature. 2012;486(7404):490-5.

67. Datar SA, Jacobs HW, de la Cruz AF, Lehner CF, Edgar BA. The Drosophila cyclin D-Cdk4 complex promotes cellular growth. EMBO J. 2000;19(17):4543–54.

68. Hennig KM, Neufeld TP. Inhibition of cellular growth and proliferation by dTOR overexpression in Drosophila. Genesis. 2002;34(1-2):107–10.

69. Stocker H, Andjelkovic M, Oldham S, Laffargue M, Wymann MP, Hemmings BA, et al. Living with lethal PIP3 levels: viability of flies lacking PTEN restored by a PH domain mutation in Akt/PKB. Science. 2002;295(5562):2088-91.

70. Lodge W, Zavortink M, Golenkina S, Froldi F, Dark C, Cheung S, et al. Tumor- derived MMPs regulate cachexia in a Drosophila cancer model. Dev Cell. 2021;56(18):2664–80 e6.

71. Poernbacher I, Crossman S, Kurth J, Nojima H, Baena-Lopez A, Alexandre C, et al. Lessons in genome engineering: opportunities, tools and pitfalls. bioRxiv. 2019:710871.

72. Hu Y, Comjean A, Perkins LA, Perrimon N, Mohr SE. GLAD: an Online Database of Gene List Annotation for Drosophila. J Genomics. 2015;3:75–81.

73. Chen EY, Tan CM, Kou Y, Duan Q, Wang Z, Meirelles GV, et al. Enrichr: interactive and collaborative HTML5 gene list enrichment analysis tool. BMC Bioinformatics. 2013;14:128.

74. Kuleshov MV, Jones MR, Rouillard AD, Fernandez NF, Duan Q, Wang Z, et al. Enrichr: a comprehensive gene set enrichment analysis web server 2016 update. Nucleic Acids Res. 2016;44(W1):W90–7.

75. Akkouche A, Mugat B, Barckmann B, Varela-Chavez C, Li B, Raffel R, et al. Piwi Is Required during Drosophila Embryogenesis to License Dual-Strand piRNA Clusters for Transposon Repression in Adult Ovaries. Mol Cell. 2017;66(3):411–9 e4.

76. Kosakamoto H, Miura M, Obata F. Epidermal tyrosine catabolism is crucial for metabolic homeostasis and survival against high-protein diets in Drosophila. Development. 2024;151(1).

77. Forero MG, Kato K, Hidalgo A. Automatic cell counting in vivo in the larval nervous system of Drosophila. J Microsc. 2012;246(2):202–12.

78. McCloy RA, Rogers S, Caldon CE, Lorca T, Castro A, Burgess A. Partial inhibition of Cdk1 in G 2 phase overrides the SAC and decouples mitotic events. Cell Cycle. 2014;13(9):1400–12.

79. Dong Q, Zavortink M, Froldi F, Golenkina S, Lam T, Cheng LY. Glial Hedgehog signalling and lipid metabolism regulate neural stem cell proliferation in Drosophila. EMBO Rep. 2021;22(5):e52130.

